# Genomic epidemiology and multilevel genome typing of Australian *Salmonella enterica* serovar Enteritidis

**DOI:** 10.1101/2022.05.18.492204

**Authors:** Lijuan Luo, Michael Payne, Qinning Wang, Sandeep Kaur, Irani U. Rathnayake, Rikki Graham, Mailie Gall, Jenny Draper, Elena Martinez, Sophie Octavia, Mark M. Tanaka, Amy V. Jennison, Vitali Sintchenko, Ruiting Lan

## Abstract

*Salmonella enterica* serovar Enteritidis is one of the leading causes of salmonellosis in Australia. However, the majority of *S*. Enteritidis cases in Australia are travel-related with a small proportion being locally acquired. This study aimed to characterise the genomic features of Australian *S*. Enteritidis and compare them with international strains using multilevel genome typing (MGT). A total of 568 *S*. Enteritidis isolates from two Australian states across two consecutive years were analysed using the *S*. Enteritidis MGT scheme and database (MGTdb) - which contained 40,390 publicly available genomes from 99 countries. The Australian *S*. Enteritidis strains were divided into three phylogenetic clades (A, B and C). Clades A and C represented 16.4% and 3.5% of the total isolates, respectively, and were of local origin. Clade B accounted for 80.1% of the isolates which belonged to seven previously defined lineages but was dominated by the global epidemic lineage (MGT4-CC1). At MGT5 level, three out of five top sequence types (STs) in Australia were also top STs in Asia, suggesting that a fair proportion of Australian *S*. Enteritidis cases may be epidemiologically linked with Asian strains. In 2018, a large egg-associated local outbreak was caused by a recently defined clade B lineage prevalent in Europe and was closely related, but not directly linked, to three isolates from Europe. Additionally, antimicrobial-resistance genes were only found in Australian clade B isolates, with a predicted multidrug resistance (MDR) rate of 11.7%. Over half (54.8%) of the MDR isolates belonged to 10 MDR-associated MGT-STs, which were also frequent in Asian *S*. Enteritidis. IncX1 plasmids were frequently present in the Australian MDR isolates. Overall, this study investigated the genomic epidemiology of *S*. Enteritidis in Australia, including the first large local outbreak, using MGT. The open MGT platform enables a standardised and sharable nomenclature that can be effectively applied to public health for unified surveillance of *S*. Enteritidis nationally and globally.

**Importance:** *Salmonella enterica* serovar Enteritidis is a leading cause of foodborne infections. We previously developed a genomic typing database – MGTdb for *S*. Enteritidis to facilitate global surveillance of this pathogen. In this study we examined the genomic features of Australian *S*. Enteritidis using the MGTdb and found that Australian *S*. Enteritidis is mainly epidemiologically linked with Asian strains (especially strains carrying antimicrobial resistance genes) followed by European strains. The first large-scale egg-associated local outbreak in Australia was caused by a recently defined lineage prevalent in Europe, and three European isolates in the MGTdb were closely related but not directly linked to this outbreak. In summary, the *S*. Enteritidis MGTdb open platform is shown to be a potential powerful tool for national and global public health surveillance of this pathogen.

## Introduction

*Salmonella enterica* serovar Enteritidis is one of the most prevalent *Salmonella* serovars causing foodborne infections [1]. *S*. Enteritidis mainly causes gastroenteritis but can also lead to non-typhoidal invasive infections (iNTS) in humans [2]. *S*. Enteritidis is widely distributed across different zoonotic niches and is especially common in poultry and poultry products like eggs [3]. The prevalence of *S*. Enteritidis increased substantially in the late 20^th^ century in Europe and North America [4]. In many countries (e.g. USA, UK, Uruguay, Lebanon), *S*. Enteritidis is ranked as the most common serovar responsible for human infections [5-8]. In the past few years, large-scale multinational outbreaks of *S*. Enteritidis were reported [9-13]. From 2017 to 2020, a multi-state outbreak of *S*. Enteritidis in Europe resulted in 656 confirmed cases [14].

Based on genome-wide comparisons, *S*. Enteritidis has been divided into three clades, clade A, B and C by Graham *et al*. [15], and five HierCC clusters by Achtman *et al* [16]. Clade B is the predominant clade which was further divided into 10 lineages in our previous study [17]. These clades and lineages have different geographical distributions [17]. Clade A and C were prevalent in Oceania but were relatively rare in other continents [17]. Within clade B, two dominant lineages represented more than 85% of clade B genomes and had different geographic distribution, outbreak propensity and evolutionary characteristics [17]. In addition to these two dominant lineages, two African lineages within clade B were associated with iNTS (mainly in sub-Saharan Africa low-income countries) and multidrug resistance (MDR) [2].

In Australia, *S*. Enteritidis is the second most prevalent serovar [18-20]. Clade A and clade C *S*. Enteritidis were previously documented in the state of Queensland (QLD) [15], while their distribution in other Australian states remained unknown. Clade B isolates in Australia were considered to be imported from other countries by travellers [15] and were rarely linked to local outbreaks [15, 21, 22]. However, a local egg-associated outbreak of clade B *S*. Enteritidis (from 2018 to 2019) caused more than 200 cases [23, 24]. In addition, antimicrobial resistance (AMR) genes were rarely present in the local clades A and C strains but were of high frequency in clade B genomes [15]. As *S*. Enteritidis has become of increasing threat in Australia, comparing the genomic features of Australian *S*. Enteritidis with international strains would help in understanding the source of Australian *S*. Enteritidis.

A unified global genomic surveillance system would contribute to tracking and controlling the spread of outbreak clusters and MDR clones in Australia. Multilevel genome typing (MGT) for *S*. Enteritidis [17] is a recently defined and publicly accessible whole genome sequencing (WGS) typing scheme (https://mgtdb.unsw.edu.au/enteritidis/). The key benefit of MGT is the provision of multiple resolution typing levels with sequence types (STs) assigned at each level [25]. The exact comparison based STs can avoid the drawbacks of clusters merging using the single linkage clustering algorithm, which enhances the stability of the ST assignments [25]. The STs from different resolution levels have been found useful in investigating the epidemiological distribution of *S*. Enteritidis and tracking MDR clones [17]. Moreover, the core genome multilocus sequence typing (cgMLST) level MGT9, which includes the serovar core genes and core intergenic regions, offered the highest resolution of typing for outbreak investigations [17, 26]. Outbreak detection clusters (ODCs) are single-linkage clusters calculated based on MGT9 allowing a different number of allele differences [25]. ODC1 (or MGT9 CC), ODC2, ODC5 and ODC10 refer to single linkage clustering with 1, 2, 5 and 10 allele differences [25]. These clustering schemes with different thresholds are built into the *S*. Enteritidis MGT database (MGTdb) and, together with metadata, allow the selection of appropriate cutoffs for optimal outbreak detection [17].

The global *S*. Enteritidis MGTdb [17] processes new publicly available genomes and metadata regularly making it useful for national and international surveillance of *S*. Enteritidis. In this study, 568 newly sequenced genomes from two states of Australia, Queensland (QLD) and New South Wales (NSW) were investigated using *S*. Enteritidis MGT. The genomic characteristics of *S*. Enteritidis from Australia and other countries were compared to assess potential Australian *S*. Enteritidis outbreak clusters and to examine the application of MGT in the national surveillance of *S*. Enteritidis using the MGTdb.

## Materials and methods

### Source of isolates

A total of 613 clinical *S*. Enteritidis isolates from QLD and NSW were collected and sequenced from 2017 to 2018 as part of routine public health laboratory surveillance. Raw reads were processed using the same methods as in our previous study [17], including contamination detection (Kraken v0.10.5) [27], quality filtering (Quast v5.0.2) [28, 29], assembling (SKESA v2.3) [30], serotyping (SISTR v1.0.2) [30], multilocus sequence typing (MLST) [1], MGT loci extraction and MGT calling [25]. A total of 568 newly sequenced *S*. Enteritidis genomes passed quality filters and were successfully processed by MGT. There were 36 (6.3%) poor quality genomes and four non-*S*. Enteritidis genomes. Isolates from the updated global MGTdb for *S*. Enteritidis included 40,390 genomes that were released to be publicly available by April 2021 (including 40,062 non-Australian and 328 Australian *S*. Enteritidis genomes excluding duplicates).

### Population structure elucidation of the Australian *S*. Enteritidis isolates

Our previous study described clades and lineages for *S*. Enteritidis using MGT STs and clonal complexes (CCs, single-linkage clustering STs with one allele difference) at different MGT levels [17]. The clades and lineages were assigned to the Australian *S*. Enteritidis genomes (896 in total) using these predefined clades/lineages markers (MGT1 to MGT5 STs/CCs) using custom Python scripts (https://github.com/Adalijuanluo/MGTSEnT).

### Identification and visualisation of clusters of closely related isolates

Closely related isolates were grouped into genomic clusters (GCs) at multiple cutoffs (i.e. 1, 2, 5 and 10 allele differences) based on the highest resolution level, MGT9. Note that GCs are equivalent to our previously defined ODCs [25]. In this study we used GCs to avoid confusion with clusters of epidemiologically confirmed outbreaks. The average number of isolates in each cluster increased as the allowed allele differences were increased from 1 to 10 (for GC1 to GC10).

From MGT9 STs to GC10 clusters, any ST or GCs with two or more isolates were identified using custom scripts. The characteristics of these clusters across the country, state and periods were depicted using Tableau v2019.2 [31]. Singletons referred to cluster types with only one isolate each in the sample. A cluster that could be potentially linked to an outbreak was defined as a non-singleton cluster with at least two isolates recovered from samples collected within a 4-week window.

Microreact [32] was used to visualise clusters at different GC levels. A dendrogram of all isolates of a GC10 cluster was constructed to visualise the division of clusters by sequentially lower GC levels. For a single GC10 cluster of interest (a GC10 cluster includes potential outbreak clusters), clusters at GC1 to 9 were recalled within the GC10 cluster. Note that the MGTdb only includes GC1 (MGT9 CC), GC2, 5 and 10. Here we also included GC3, 4, 6, 7, 8 and 9, to increase the flexibility of outbreak clusters selection. A dendrogram was produced based on the recalled cluster types from GC1 to GC10, as well as MGT9 STs. The root was the GC10 cluster, each leaf was an MGT9 ST and each internal node represented a GC cluster between GC1 and GC9. Therefore, the dendrogram is a GC typing tree rather than a phylogenetic tree. The dendrogram and the metadata were visualised in Microreact [32]. Isolates with invasive infections can be highlighted in Microreact [32]. The geographic distribution and isolation timeline of different subclusters can be visualised and compared in Microreact [32].

### Bayesian evolutionary analysis of the Australian outbreak lineage

Bayesian evolutionary analysis was performed to investigate the evolutionary history of the isolates associated with the Australian outbreak. For the isolates belonging to the European common lineage, 144 isolates have unique MGT8 STs which were sampled for phylogenetic tree construction using quicktree.pl (which is a pipeline included in SaRTree v1.2.2) with P125109 as reference [33]. Ten sampled Australian outbreak isolates were phylogenetically close to 18 international isolates and one non-outbreak Australian isolate, which belonged to the same branch. These 29 isolates were selected to estimate the evolutionary history of the large-scale egg-associated outbreak in Australia, Bayesian evolutionary analysis was performed using BEAUti v1.10.4 and BEAST v1.10.4 [34]. Eight models were evaluated with GTR gamma model, MCMC chain length of 100,000,000, and the phylogenetic expansion effective sample size (ESS) of > 200 [34]. The population expansion was estimated using Tracer.v1.7.1 with 10% burnin [35]. SNP alignments were produced from the MGT9 allele profile. RecDetect v1.2.2 [33] was used to remove the recombination sites on the SNP alignments. A maximum-likelihood tree (GTR, gamma model) was constructed using MEGA X [36] with a bootstrap number of 1000. The log file of the Bayesian skyline model was mapped onto the maximum likelihood tree using BEAST tree annotator v1.10.4 and visualized using Figtree v1.4.4 [34, 37].

### AMR prediction and plasmid replicon genes analysis

AMR associated genes and mutations were screened using the AMRFinderPlus v3.10.18 database using the default settings [38, 39]. Our previous study [17] estimated the presence of AMR genes in all publicly available genomes of *S*. Enteritidis and identified MGT-STs that were associated with MDR using custom python scripts. These MDR associated STs were identified among the Australian *S*. Enteritidis genomes. In addition, plasmid replicon genes were screened against the PlasmidFinder v2.1 database [40] through Abricate v1.0.1 (https://github.com/tseemann/abricate). Plasmid replicon genes with identity and coverage >= 90% were regarded as positive.

## Results

### Epidemiological characteristics of the sampled isolates

A total of 896 Australian *S*. Enteritidis isolates were analysed, of which 568 were newly sequenced genomes from two Australian states, QLD (257) and NSW (311), and the remainder (328) were publicly available genomes (**Table 1**). The newly sequenced isolates were collected between June 2017 to November 2018 (**Figure 1**), except for one isolate which was collected in July 2019. The collection period of all Australian isolates (including publicly available ones) was from 2001 to July 2019.

**Table 1.** *S*. Enteritidis genome sequences used in this study.

|  | Australia |  |  | Other<br>countries | Total |
| --- | --- | --- | --- | --- | --- |
|  | QLD | NSW | Other states |  |  |
| Newly sequenced | 257 | 311 |  |  | 568 |
| Publicly available | 152 | 1 | 175 | 40,062 | 40,390 |
| Total | 409 | 312 | 175 | 40,062 | 40,958 |

**Figure 1.**
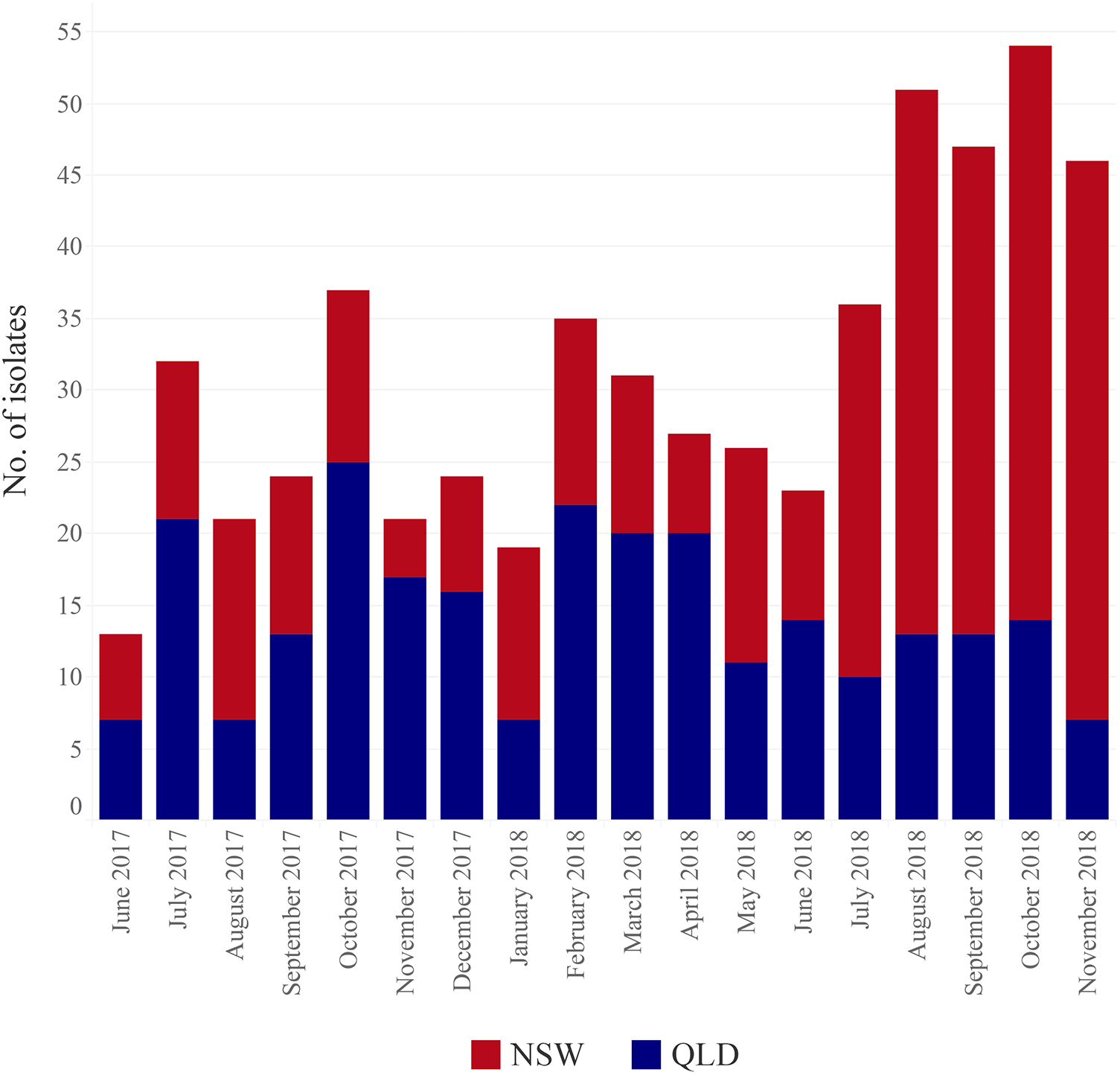
The collection dates of the newly sequenced genomes from NSW and QLD in Australia. The X-axis shows the month and the year of the collection time while Y-axis shows the number of isolates for each month.

The newly sequenced Australian *S*. Enteritidis genomes were compared to the updated *S*. Enteritidis MGTdb with 40,390 publicly available genomes which were collected from 99 different countries from 1917 to 2021. The detailed information of these genomes is described in the **supplementary text, Figure S1** and **Table S1**.

### MGT typing of newly sequenced isolates

The 568 newly sequenced Australian *S*. Enteritidis genomes were processed through the *S*. Enteritidis MGTdb and held as private data. STs were called from MGT2 to MGT9. MGT1 refers to the classic seven-gene MLST [1], which typed 99.7% of the 568 genomes into five STs. ST11 (78.9%) was the predominant type, followed by ST3304 (9.5%), ST1925 (7.0%), ST180 (5.3%) and ST1972 (1.9%). With the increasing resolution of typing from MGT2 to MGT9, there was an increasing number of STs called from 23 STs at MGT2 to 482 at MGT9 (**Figure 2a**). Note that four genomes (0.7%, 4/568) failed to be called at the MGT9 level due to 2.2% to 2.6% of missing alleles.

**Figure 2.**
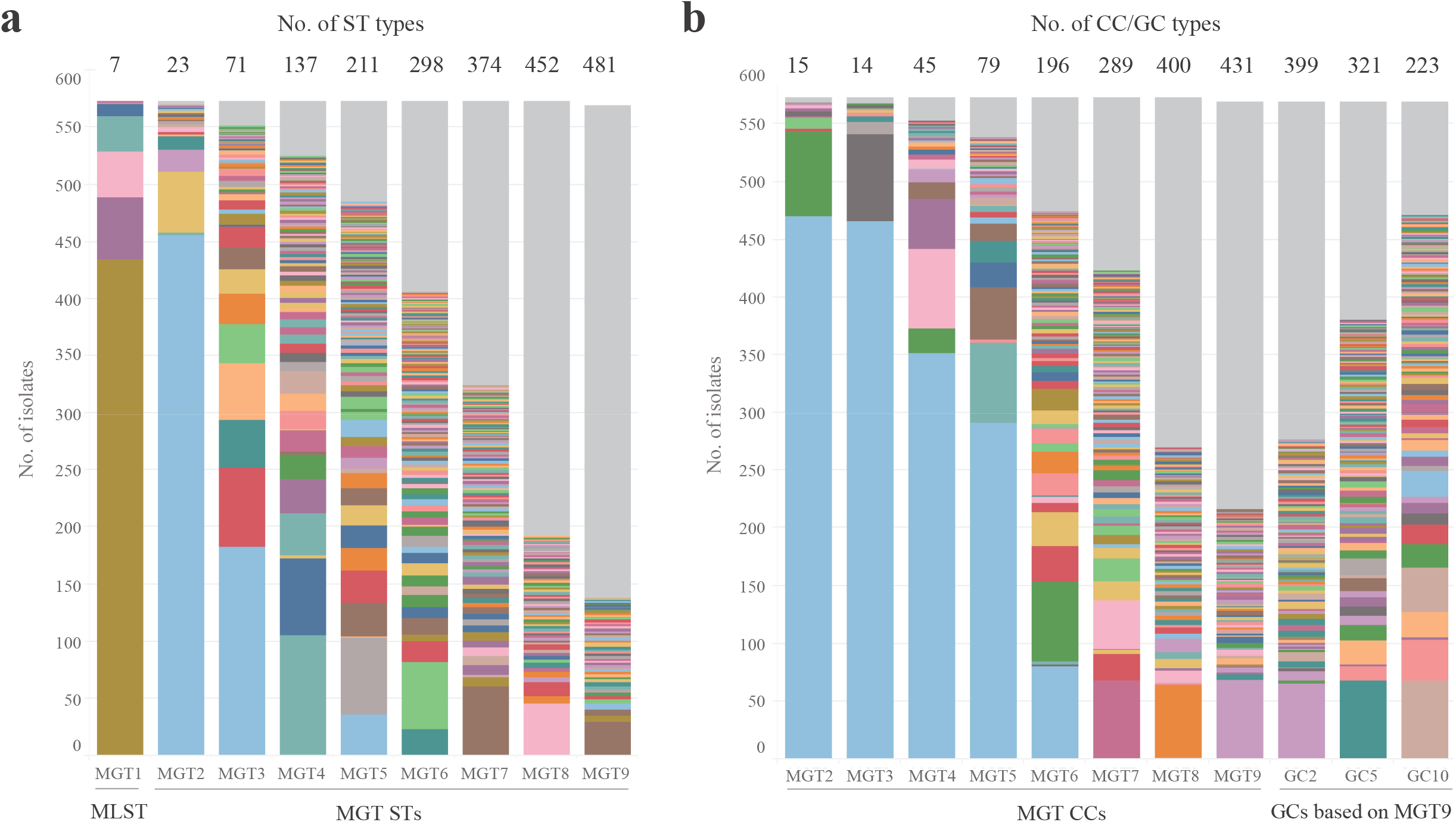
MGT typing of the newly sequenced genomes. **a**. Sequence types (STs) assigned at each MGT level. **b**. Single linkage clustering of STs into clonal complexes (CCs) (assigned based on STs from MGT2 to MGT9) and genomic clusters (GCs) (assigned based on MGT9-STs). STs/CCs/GCs >= 2 isolates each were separated into different colours and singleton types (one isolate each) were grouped in grey.

From MGT2 to MGT9, STs were grouped into CCs using the single linkage clustering algorithm with one allele difference as cutoff (**Figure 2b**), using the same approach and definition as classic MLST [41]. MGT9 STs were further grouped using the single linkage clustering algorithm with more allele differences allowed (i.e. 2, 5 and 10 allele differences, which were used to generate GC2, GC5 and GC10, respectively) as described previously (**Figure 2b**) [17, 25].

### Population structure of Australia *S*. Enteritidis isolates by clades and lineages

The population structure of *S*. Enteritidis was previously defined with three clades (A, B and C) and 10 lineages within clade B [15, 17]. Australian isolates fell into these clades and lineages. By clades, 16.4% (147), 80.1% (718) and 3.5% (31) of the 896 Australian *S*. Enteritidis genomes belonged to clade A, B and C, respectively (**Figure 3**). Within clade B, seven of the previously defined 10 lineages were observed in Australia, representing 97.5% (700/718) of the clade B genomes (**Figure 3**). The global lineage MGT4-CC1 was the predominant lineage (81.8%, 587/718), followed by the European lineage MGT4-CC30 (10.0%, 72/718) (**Figure 3**). MGT4-CC13, which is especially prevalent in North America and Europe [17], was relatively rare in Australia (4.6%, 33/718). Three isolates belonged to the Africa endemic iNTS associated lineages MGT3-CC10 (two MGT3-ST365 strains) and CC15 (one MGT3-ST15) and were cultured from blood, wound and ankle fluid samples, respectively. Only 18 clade B isolates (2.5%, 18/718) belonged to none of the 10 reported lineages.

**Figure 3.**
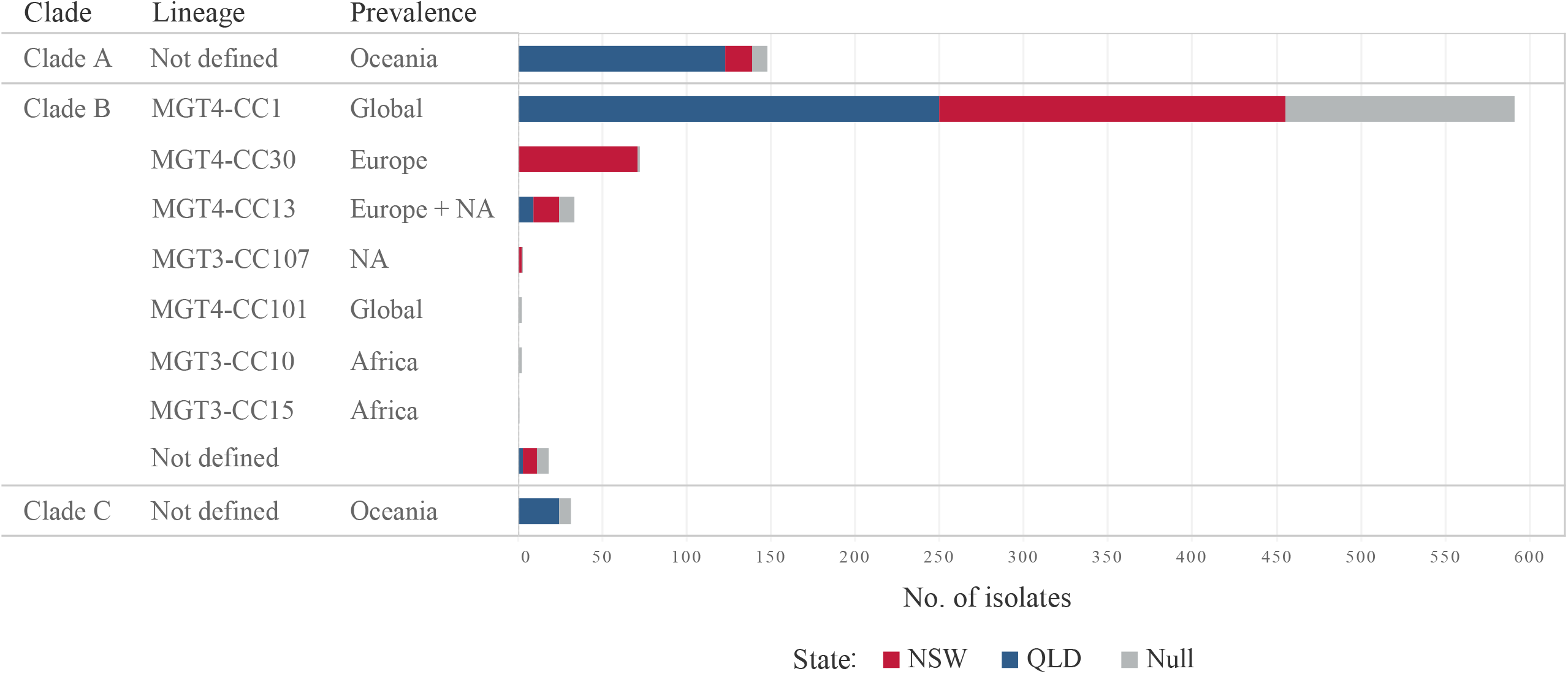
The population structure of the Australian *S*. Enteritidis using existing nomenclatures. NA = North America. The population structure of all Australian isolates (896) including the publicly available genomes. Seven previously defined lineages of clade B were observed, with the majority of isolates belonging to the global lineage MGT4-CC1. The two African lineages were observed with three isolates. Note the geographic prevalence of different clades and lineages was based on the metadata in MGTdb which may have sampling bias.

### Comparison of Australian isolates across states and with international isolates using MGT5

Our previous study found MGT5 was the optimal resolution level to depict the national genomic epidemiology of *S*. Enteritidis [17]. Hence, in the current study, the geographic and temporal genomic characteristics of the newly sequenced Australian *S*. Enteritidis genomes were analysed at the MGT5-level. We observed that there were 19 MGT5-STs with >= 5 isolates each, representing 55.6% of the genomes (**Figure 4**). For clade C, the 11 isolates were sporadically observed in seven months with no STs of more than five isolates each.

**Figure 4.**
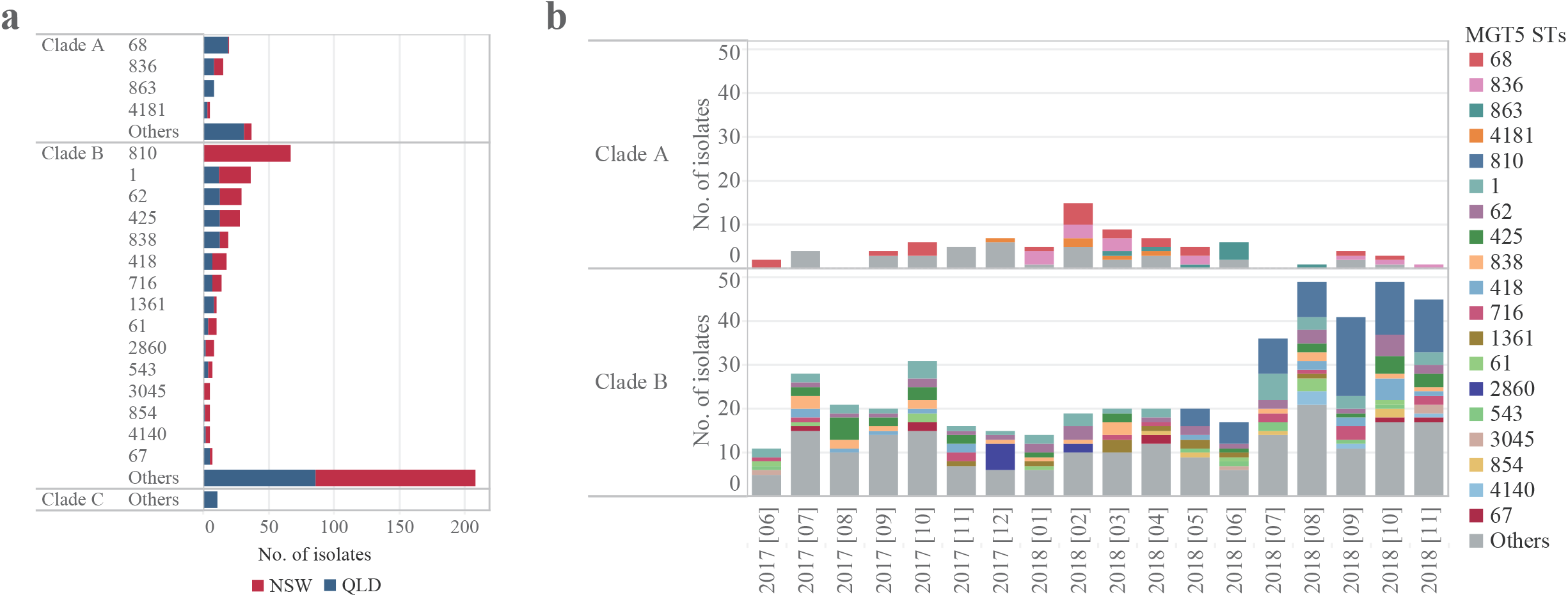
Genomic epidemiological characteristics of the MGT5-STs in Australia. **a**. The main MGT5-STs (>=5 isolates each) of the newly sequenced Australian *S*. Enteritidis genomes, and their distribution by states (NSW and QLD). **b**. Distribution of the main MGT5-STs in different months. STs with >= 5 isolates each were represented with different colours. Some STs occurred almost every month whereas several STs appeared and persisted over several months.

In clade A, there were four MGT5-STs with >= 5 isolates each, all of which were only observed in Australia. Three STs were observed in both QLD and NSW (MGT5-ST68, ST836, ST4181), and one ST (MGT5-ST863) was only observed in QLD (**Figure 4a**). Across the 18-month sampling window from June 2017 to October 2018, MGT5-ST68 was observed over 10 months, MGT5-ST836 and MGT5-ST863 were observed over seven and five months respectively, while other STs were observed over shorter periods (**Figure 4b**). The top three STs, MGT5-ST68, ST836 and ST863, were found in the publicly available genomes from Australia in the MGTdb, whereas ST4181 was a newly identified type **(Table 2**). MGT5-ST68 ranked as the top ST and was first observed in 2005 in QLD while the other three STs were observed from 2016 onwards (**Table 2**).

**Table 2.** Comparison of the newly sequence genomes with genomes in the S. Enteritidis MGTdb by MGT5-STs.

| S. Enteritidis MGTdb <sup>a</sup> |  |  |  |  |  |  |  |  |  |  |  |  |  |  |  |
| --- | --- | --- | --- | --- | --- | --- | --- | --- | --- | --- | --- | --- | --- | --- | --- |
| MGT5 STs | Clades | MLST STs | Clade B MGT4 | No. of new isolates | No. of total isolates | Oceania <sup>b</sup> | Asia | Europe | North America | South America | Africa | No. of countries | Top country | Earliest Year | Invasive |
| 68 | A | 3304 |  | 20 | 38 | Top 4 |  |  |  |  |  | 1 |  | 2005 |  |
| 836 | A | 180 |  | 14 | 17 | + |  |  |  |  |  | 1 |  | 2016 |  |
| 863 | A | 3304 |  | 8 | 9 | + |  |  |  |  |  | 1 |  | 2017 |  |
| 4181 | A | 3304 |  | 5 | 5 |  |  |  |  |  |  | 1 |  | 2017 |  |
| 810 | B | 11 | CC30 | 67 | 97 | + |  | + | + |  |  | 7 | Aus | 1981 |  |
| 1 | B | 11 | CC1 | 36 | 2353 | Top 1 | Top 1 | Top 1 | Top 17 | Top 1 | Top 2 | 39 | UK | 1981 | 2.5% |
| 62 | B | 11 | CC1 | 30 | 92 | Top 2 |  | + | + |  |  | 4 | Aus | 2013 |  |
| 425 | B | 11/1925 | CC1 | 28 | 167 | Top 3 | Top 5 | + | + |  |  | 8 | UK | 2008 | 1.2% |
| 838 | B | 11 | CC1 | 19 | 53 | + |  | + | + |  |  | 4 | UK | 2015 |  |
| 418 | B | 11 | CC1 | 18 | 64 | Top 5 | + | + | + |  |  | 4 | Aus | 2015 |  |
| 716 | B | 11 | CC1 | 14 | 65 |  |  | + | + |  |  | 5 | UK | 2016 |  |
| 1361 | B | 11 | CC1 | 10 | 17 | + |  | + | + |  |  | 3 | Aus | 2017 |  |
| 61 | B | 1925 | CC1 | 10 | 98 | Top 6 | Top 3 | + | + |  |  | 5 | UK | 2012 | 1.0% |
| 2860 | B | 1925 | CC1 | 8 | 10 |  |  | + |  |  |  | 2 | Aus | 2017 |  |
| 543 | B | 11 | CC1 | 7 | 56 | + | + | + | + |  |  | 5 | UK | 2014 | 3.6% |
| 3045 | B | 11 | CC1 | 5 | 7 |  |  | + |  |  |  | 2 | Aus | 2017 |  |
| 854 | B | 11 | CC1 | 5 | 12 | + |  | + |  |  |  | 2 | Aus/UK | 2016 |  |
| 4140 | B | 11 | CC1 | 5 | 5 |  |  |  |  |  |  | 1 |  | 2018 |  |
| 67 | B | 11 | CC13 | 7 | 201 | + |  | Top 19 | + |  | Top 4 | 8 | UK | 2004 | 6.5% |

<sup>a</sup> Top 20 STs were labelled, only STs >= 5 isolates each were included for ranking. + represented the continent containing the ST but it ranked outside of the top 20 STs.
<sup>b</sup> Publicly available genomes from Oceania with 99% from Australia.

In clade B, there were 15 MGT5-STs with >= 5 isolates, 86.7% (13/15) of the STs were found in both states (**Figure 4a**). The 473 newly sequenced Australian clade B genomes were collected over 18 months (**Figure 4b**). MGT5-ST1, ST62, ST425, ST838 and ST418 occurred almost every month. However, several STs were not continuously observed but instead appeared and persisted over several months. These STs included MGT5-ST810 emerged in May 2018, MGT5-ST2860 emerged in December 2017. Both STs were observed with 5 or more isolates in one month.

By comparing the 15 main MGT5-STs of clade B in Australia with the publicly available genomes in the MGTdb, 14 (93.3%, 14/15) STs were also observed in other countries (**Table 2**), the dominant country of which was either Australia or the UK. MGT5-ST1 is the most frequently ST observed in multiple continents, including Oceania, Asia, Europe, South America and the second most frequent ST in Africa. Despite limited sampling from Asia (693 genomes in total in the MGTdb), five Australian STs were found in Asia, with three STs being ranked as the top, second and fifth most common STs in Asian isolates (**Table 2**). In contrast, the majority of the clade B STs that were also found in Europe and North America were rarely ranked as the top STs in these places (except for MGT5-ST1). In terms of collection time, MGT5-ST810 and MGT5-ST1 were first sampled in 1981, whilst the other STs were first sampled in 2013. In addition, five Australian STs contained isolates that were collected from blood or cerebrospinal fluid (1.0% to 6.5% of the isolates for each ST) suggesting that these STs have caused invasive human infections.

### Using a 4-week window to identify potential outbreak clusters by different resolution levels of GCs

MGT9, which included 4986 loci, offered the highest resolution to identify closely related isolates. We further used GCs which are single-linkage clusters based on the MGT9 alleles to examine closely related clusters of isolates that were found across states and countries. GC1 (i.e. MGT9 CC), GC2, GC5 and GC10 corresponded to allele difference cutoffs of 1, 2, 5 and 10 respectively for clustering MGT9 STs. Note that GCs are purely an alternative method of grouping isolates at small genetic distances. Using GCs solely without any other metadata cannot define outbreaks. Using the MGTdb, we compared the GC clusters across countries and states (NSW and QLD) as introduced in the **supplementary text** and **Figure S2 to S3**.

For *Salmonella* outbreak investigation in NSW using multilocus variable-number tandem repeat analysis (MLVA), Sintchenko *et al*. defined a cluster of 5 cases with a specific MLVA type within a 4-week window as a threshold signalling an outbreak [42]. However, for WGS data, no threshold of genome differences or a minimum number of cases has been defined. We used a 4-week window and a minimum of 2 cases as a putative cluster definition of relevance to public health surveillance in this study. At different GC levels, the GC clusters that contained at least two isolates within 4 weeks (i.e. the minimum pairwise collection date distance of isolates <= 4 weeks) were illustrated (**Figure 5**). While the clusters identified may also include isolates more than 4 weeks apart. Clusters with no collection date information were grouped as unknown. No epidemiological investigation metadata for the newly sequenced genomes was available to confirm whether any isolates were outbreak-related except for the large outbreak in NSW described below.

**Figure 5.**
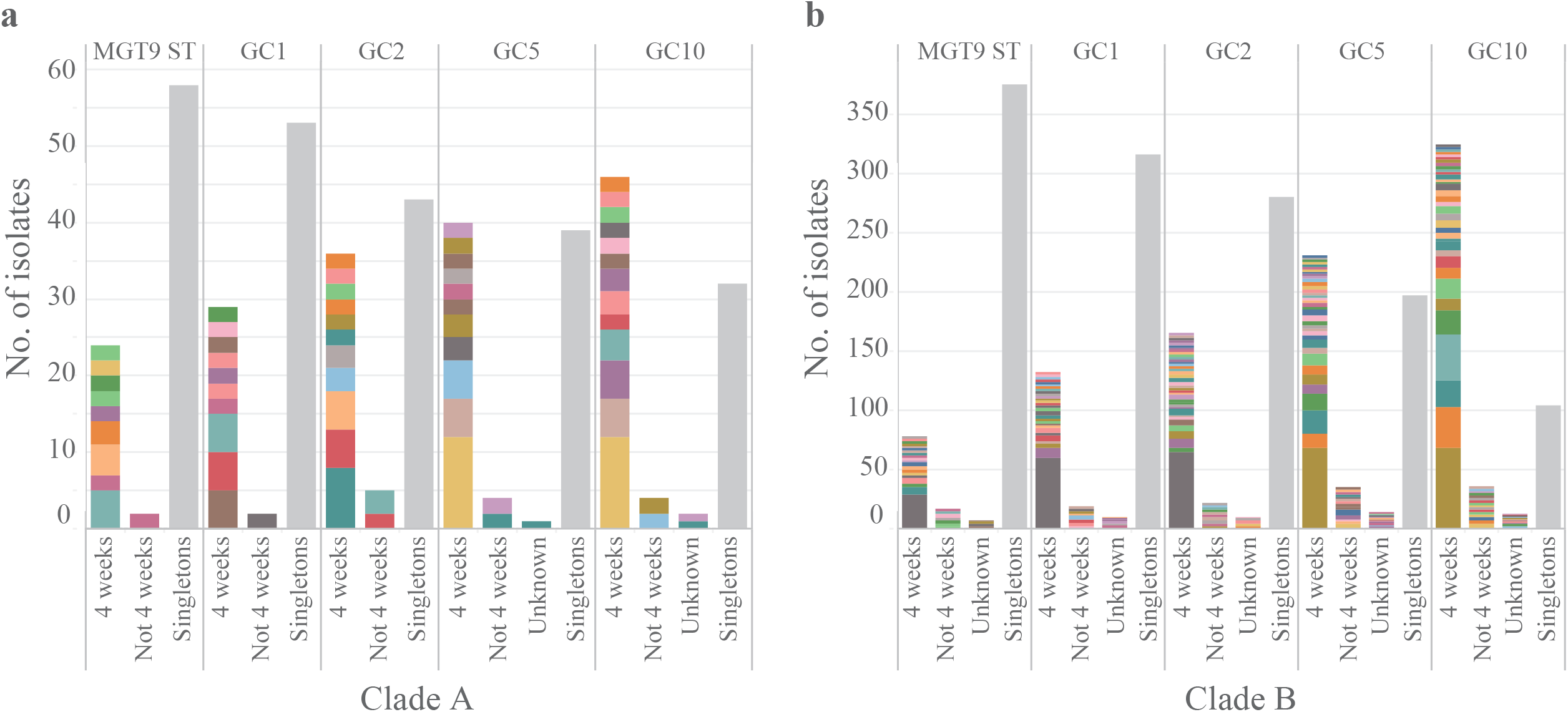
Identification of putative outbreak clusters using a 4-week window. Clusters with at least two isolates occurring within four weeks in clade A and B were identified at different resolution genomic cluster (GC) levels. Clusters at each GC level were categorised into four groups. Singletons were clusters with only one isolate each, which were collapsed into one column in grey colour. For the non-singleton clusters: “4 weeks” referred to the clusters including at least two isolates collected within 4 weeks; “Not 4 weeks” referred to clusters that did not include two isolates collected within 4 weeks; ‘Unknown’ referred to clusters without collection date information.

For newly sequenced clade A genomes (84 isolates), there were 9, 10, 11, 11 and 13 clusters including isolates collected within 4 weeks at MGT9-ST, GC1, 2, 5 and 10, respectively (**Figure 5a**). These clusters represented 28.6%, 34.5%, 42.9%, 46.4% and 53.6% of the total 84 newly sequenced clade A genomes at MGT9-ST, GC1, 2, 5 and 10, respectively. For clade B, there were 20, 28, 38, 37 and 35 clusters including isolates collected within a 4-week window at MGT9-ST, GC1, 2, 5 and 10, respectively (**Figure 5b**). These clusters accounted for 16.5%, 27.5%, 34.7%, 48.2% and 67.9% of the 473 newly sequenced clade B genomes (from NSW and QLD) at MGT9-ST, GC1, 2, 5 and 10, respectively. For clade C, only one cluster was collected in 4 weeks at GC5 and GC10.

### Evolutionary analysis of the first Australian clade B *S*. Enteritidis outbreak of local origin

A confirmed large-scale outbreak of clade B *S*. Enteritidis occurred in Australia in 2018 and lasted until 2019 and was the first known *S*. Enteritidis outbreak to have originated in a local farm source [23, 24]. This outbreak was associated with contaminated eggs in NSW and caused more than 200 confirmed cases [23, 24]. The majority of the cases were in NSW, but some were also observed in other states of Australia, and New Zealand [23, 24]. The present study analysed 68 genomes of outbreak isolates from NSW collected in 2018.

Different GC levels were examined to identify the level that best described the outbreak. By GC10, one cluster contained all Australian outbreak isolates but also contained 10 international isolates. By GC5, the number of international isolates was reduced to three with two isolates from the UK and one from Ireland isolates which were isolated from 2012 to 2015. By GC4, only the NSW isolates were clustered together without any international isolates. The NSW isolates mainly belonged to MGT7-ST3370 (90.0%).

To determine the evolutionary origins of the local *S*. Enteritidis lineage, we randomly sampled 29 genomes from the European common lineage MGT4-CC30 to which the Australian outbreak lineage belonged. Exponential growth and strict molecular clock were evaluated as the optimal combination of settings in BEAST with a mean ESS greater than 200. The most recent common ancestor (MRCA) of the Australian outbreak isolates was estimated to be around 2015 (95% CI 2013-2017) (**Figure 6**, green background). The MRCA of the Australian outbreak isolates and the UK isolates, which were the most closely related sampled isolates with the Australian outbreak, was estimated to be around 2011 (95% CI 2008-2012) (**Figure 6**). The Bayesian phylodynamic and population expansion of this lineage is shown in **Figure S4**.

**Figure 6.**
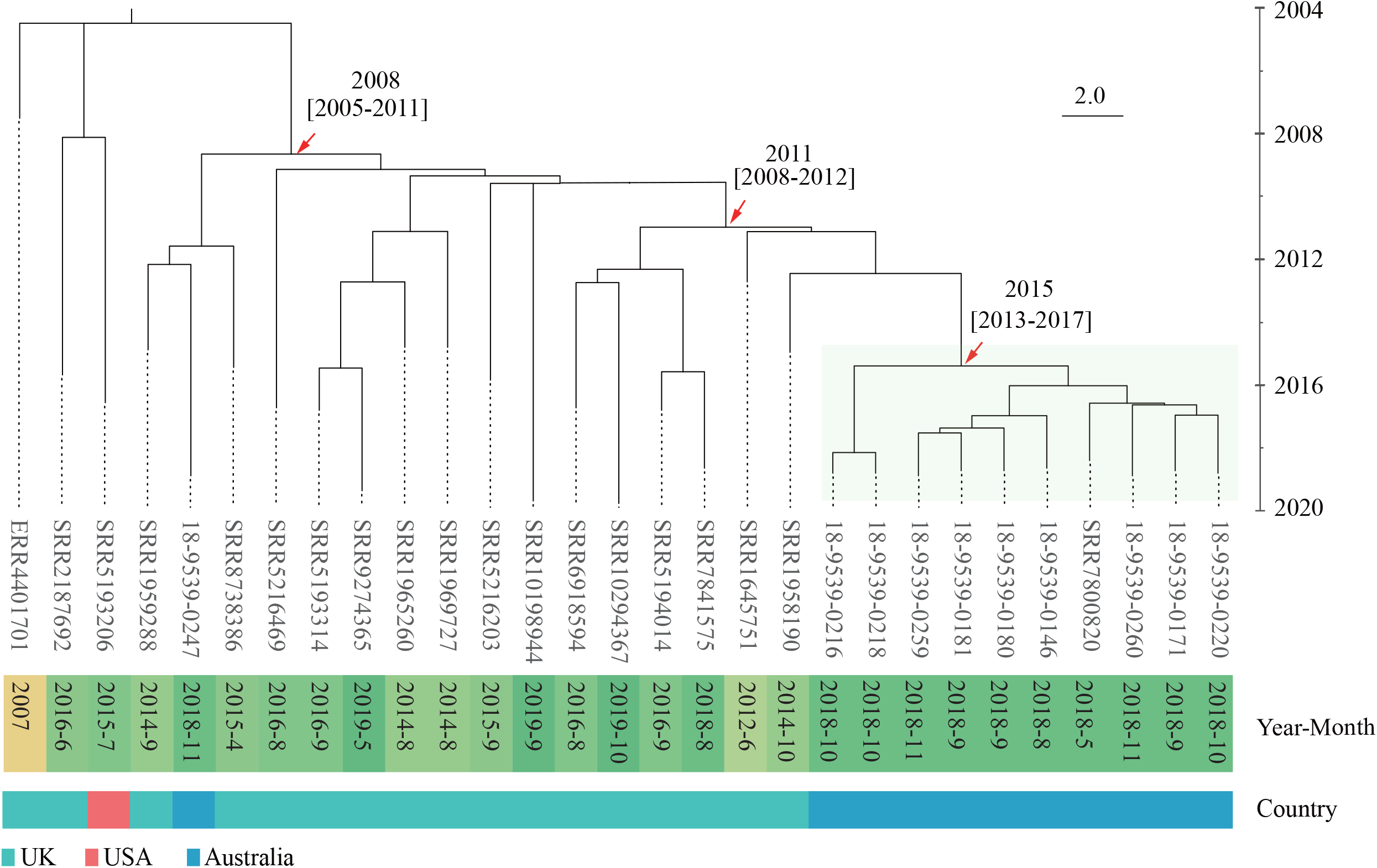
Phylodynamic analysis of the NSW outbreak related isolates. A total of 29 isolates were sampled with the exponential growth strict clock model as the optimal model. Country and collection year-month information of the sampled isolates were represented with different colours. Isolates belonging to the Australian outbreak were highlighted in green. The most recent common ancestor (MRCA) of the Australian outbreak isolates was around 2015 (95% CI 2013-2017). Six isolates from the UK were phylogenetically close to the Australian outbreak isolates, and the MRCA was around 2011 (95% CI 2008-2012). Note that one Australian isolate from 2018 that was unrelated to the large outbreak was identified and included in the tree.

### Using Microreact to visualise and investigate large international outbreak clusters spread to Australia

International outbreak-related clusters have been observed in Australia. To visualise the temporal and spatial distribution of a single international cluster that includes Australian isolates, Microreact [32] in combination with a GC dendrogram were used. To investigate the relationship between the Australian isolates and international isolates and to demonstrate the usefulness of this approach, one large international cluster GC10-Cluster 89 (C89) was examined (**Figure 7a)**. GC10-C89, which was observed from 2004 to 2021, included 392 isolates from nine countries, including 11 isolates from Australia. Six of the 11 Australian isolates belonged to three GC2 clusters (two isolates each) being collected within a 4-week window, suggesting potential small outbreak clusters. The leaves of the dendrogram represented isolates with different MGT9-STs, nodes represented subclusters from GC1 to GC9 within GC10-C89, which can be selected and visualised as a subtree (**Figure 7a)**. An international subcluster with 24 isolates was defined by a GC4 cluster (GC4-C5) (**Figure 7b**). This GC4-C5 cluster was observed in Australia, the UK, the USA, South Africa and Ireland from 2004 to 2018. Breaking down further to higher resolution GC levels, two Australian *S*. Enteritidis isolates, which were collected in April 2018, were of the same GC2 type as the three isolates from the UK, USA and Ireland, two of which were also found in 2018 (unknown year for the USA isolate). The data suggest a potential international outbreak cluster transmitted to Australia. Other isolates with different GC2 cluster types were collected from 2004 to 2017. Notably, according to the publicly available metadata of source types, GC10-C89 included 24 invasive infection isolates which were observed in South Africa, Australia and the USA (**Figure 7**). The interactive version of the GC10-C89 dendrogram and the linked spatial, temporal and metadata information of **Figure 7** are available online in Microreact (https://microreact.org/project/6y1Qay9BRdvZBdSGTWzfHr).

**Figure 7.**
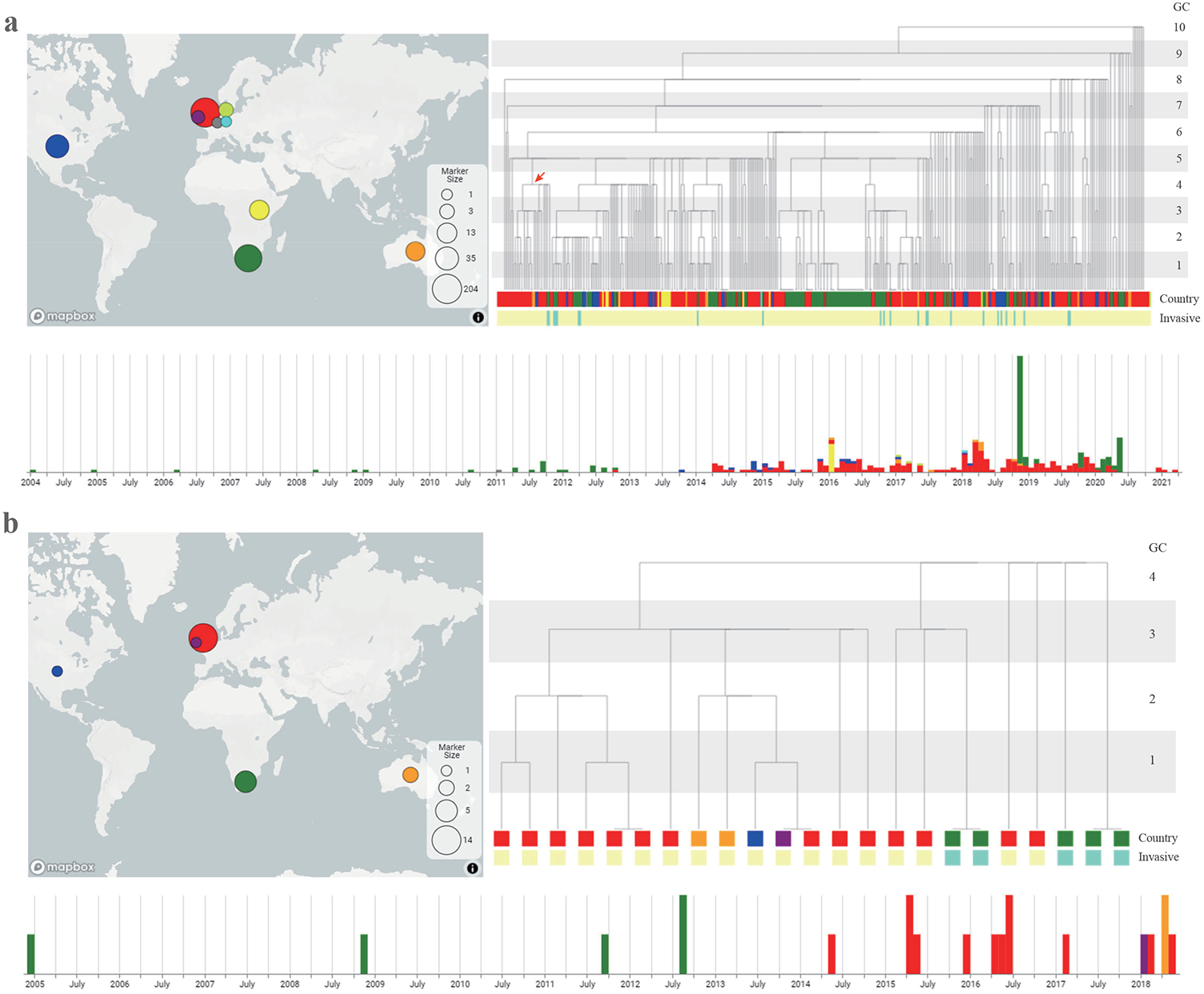
Visualisation of an international cluster (GC10-C89) in Microreact. **a**. Visualisation of the whole cluster. The dendrogram was generated based on the cluster types from different resolution levels (MGT9 STs to GC10). The root was the GC10 cluster and each leaf was an MGT9 ST. Each internal node was a GC between GC1 and GC9 as labelled. At each GC level, the branches that were linked with a horizontal line belonged to the same cluster type. Different colours were used to represent different countries. Isolates caused possible invasive infections were labelled in green colour band. The size of the circles in the map corresponds to the number of isolates. Note that international isolates with collection year and month information but no date were assumed to be collected on the 15th of each month. Subclusters at different GC levels can be selected and visualised. **b**. Visualisation of a GC4 subcluster (marked by the red arrow in a) as a separate tree. The two Australian *S*. Enteritidis genomes collected from 2018 were of the same GC2 type with three isolates from the UK, USA and Ireland, respectively. The UK and Ireland isolates were also observed in 2018.

### Presence of AMR genes in Australian *S*. Enteritidis

Antimicrobial resistance is emerging in *S*. Enteritidis, especially in the two lineages prevalent in Africa [2]. We evaluated the presence of AMR genes in the Australian *S*. Enteritidis genomes. No AMR genes were observed in isolates from clade A and C. In clade B, 69.6% (500/718) were found to possess AMR genes, 11.7% (84/718) were predicted to be MDR, with genes conferring resistance to three or more drug classes (**Table 3**). Among the newly sequenced isolates, 11.6% of clade B were predicted to be MDR. By lineages, 97.6% (82/84) of the MDR isolates belonged to the global lineage MGT4-CC1.

**Table 3.**
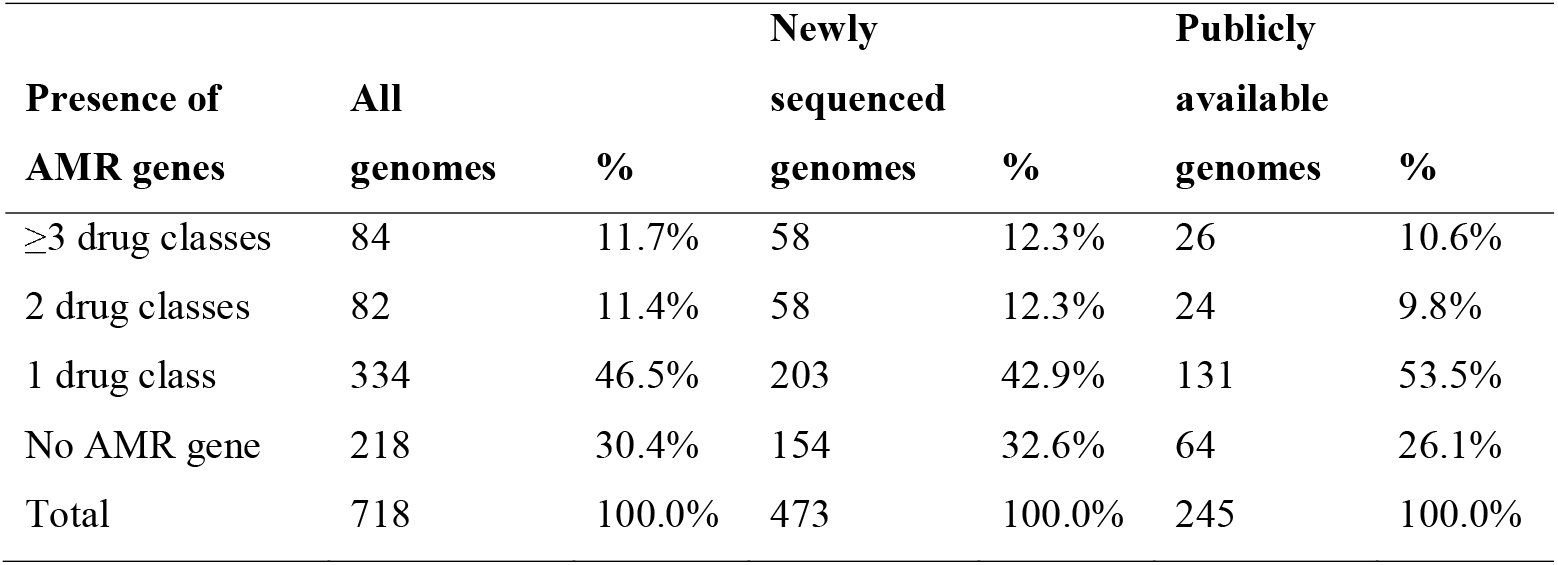
Presence of antimicrobial resistance (AMR) genes in Australian clade B genomes.

| **Presence of**<br>**AMR genes** | **All** |  | **Newly**<br>**sequenced** |  | **Publicly**<br>**available** |  |
| --- | --- | --- | --- | --- | --- | --- |
|  | **genomes** | **%** | **genomes** | **%** | **genomes** | **%** |
| $\geq 3$ drug classes | 84 | 11.7% | 58 | 12.3% | 26 | 10.6% |
| 2 drug classes | 82 | 11.4% | 58 | 12.3% | 24 | 9.8% |
| 1 drug class | 334 | 46.5% | 203 | 42.9% | 131 | 53.5% |
| No AMR gene | 218 | 30.4% | 154 | 32.6% | 64 | 26.1% |
| Total | 718 | 100.0% | 473 | 100.0% | 245 | 100.0% |

We previously identified MDR associated STs within which there was a high proportion of predicted MDR genomes (>= 80%) at different MGT levels [17]. All Australian *S*. Enteritidis genomes (896 isolates) were screened for these MDR associated STs. There were 47 Australian *S*. Enteritidis isolates found in 10 MDR associated MGT-STs (**Table 4**) and 97.9% (46/47) of these isolates were confirmed to carry MDR genes. Therefore, among the 84 Australian clade B genomes carrying MDR genes, 54.8% (46/84) belonged to previously identified MDR associated MGT-STs. The AMR drug classes of the 10 MDR associated STs were shown in **Table 4**, MGT4-ST54, MGT6-ST107 and MGT4-ST354 were the top MDR associated STs in Australian *S*. Enteritidis.

**Table 4.** Multidrug resistance (MDR) associated MGT-STs in Australian *S*. Enteritidis and the drug classes.

| MGT-ST | Lineage | No. of<br>Australian<br>isolates | Drug classes <sup>a</sup> |  |  |  |  |  |  |
| --- | --- | --- | --- | --- | --- | --- | --- | --- | --- |
|  |  |  | Aminoglycoside | Beta-lactam | Phenicol | Quinolone | Sulphonamide | Tetracycline | Trimethoprim |
| MGT3-ST15 | MGT3-CC15 | 1 | + | + | + |  | + | + | + |
| MGT4-ST54 | MGT4-CC1 | 15 | + | + |  | +/- | + | + |  |
| MGT6-ST107 | MGT4-CC1 | 10 | + | + |  | + | + |  |  |
| MGT4-ST354 | MGT4-CC1 | 8 | + | + |  | + | + | +/- |  |
| MGT3-ST9 | MGT4-CC1 | 4 | + | + |  | +/- | + | + |  |
| MGT3-ST435 | MGT4-CC1 | 3 | + | + |  | + | + | +/- |  |
| MGT5-ST86 | MGT4-CC1 | 3 | + | + |  | + | + |  |  |
| MGT4-ST955 | MGT4-CC1 | 1 | + | + |  | + | + |  |  |
| MGT4-ST1419 | MGT4-CC1 | 1 | + | + |  | + | + |  |  |
| MGT5-ST2729 | MGT4-CC1 | 1 | + | + |  | + | + |  |  |

<sup>a</sup> The drug classes were predicted by ResFinder [61]. + means $\geq 80\%$ of the isolates were predicted to be resistant to the drug class; +/- means $< 80\%$ but $\geq 20\%$ of isolates were predicted to be resistant to the drug class.

Plasmids were associated with the acquisition of AMR genes. We screened the Australian *S*. Enteritidis genomes for plasmid replicon types. In clade A, only 10.9% (16/147) of isolates had plasmid replicon genes (10 types), with IncI1 as the most common type observed in eight isolates (**Table S2**). In clade C, 4 out of 31 isolates (12.9%) had plasmid replicon genes observed belonging to four different types. In clade B, 32 plasmid replicon types were found in one or more clade B genomes with 97.5% (700/718) isolates carrying at least one replicon gene (**Table S2**). Except for IncFII(S) and IncFIB(S) which were present in > 94% of the clade B isolates, InxX1_4 was the dominant type (22.1%), followed by IncX1 (12.1%) and Col156 (11.0%). We further estimated the presence of AMR genes in isolates possessing different plasmid replicon types (**Figure S5**). Except for IncFII(S) and IncFIB(S), more than 85% of the isolates with IncX1 (IncX1_1 or IncX1_4) plasmid types were found to carry AMR genes (**Figure S5**). Nearly 40% of isolates with IncX1_4 plasmids were predicted to be MDR.

## Discussion

In Australia, *S*. Enteritidis is the second most common serovar after *S*. Typhimurium causing human non-typhoidal salmonellosis, however, *S*. Enteritidis appears to be not prevalent in Australian farms [18-20]. Most *S*. Enteritidis infections have been associated with travel and there were relatively few locally acquired cases [18-20, 43]. Therefore, the molecular epidemiology of *S*. Enteritidis in Australia has been significantly different from locally prevalent *Salmonella enterica* serovars and distinct from other countries. To investigate the genomic epidemiology of Australian *S*. Enteritidis, this study characterised 896 Australian genomes (568 were newly sequenced), by comparing them with ∼40,000 international genomes in the MGTdb. All Australian *S*. Enteritidis isolates belonged to clade A, B or C, with clade B being dominant (80.1%), consistent with a previous report [15]. An additional two small clades were recently identified from global data [16], but neither was found in Australian *S*. Enteritidis.

### Australian clades A and C are rarely linked to international isolates and clade A can cause small-scale local outbreaks

In the global *S*. Enteritidis MGTdb, more than 85% of clade A and 70% of clade C genomes were from Australia. Using seven gene MLST (MGT1), clade A includes two STs and clade C only one. MGT1-ST3304 (clade A) was not observed in any other country, as was previously reported [15]. MGT1-ST180 (clade A) was also found in New Zealand, Vanuatu, Vietnam, Germany, the UK and the USA. MGT1-ST1972 (clade C) was also found in the UK and the USA. Within Australia, clade A was significantly more common in QLD than in NSW and clade C was only observed in QLD. This agrees with a previous report that QLD had the largest number of locally acquired *S*. Enteritidis infections [19]. Factors that affect the differences in the distribution of the two clades between states in Australia are unknown.

Using GCs and a 4-week window for clustered isolates as potential outbreaks, the data showed a few small clusters that were potentially outbreak clusters, although there was no epidemiological information to confirm them. For clade A, there were multiple small GC2 clusters with 2-8 isolates each within clade A that included isolates collected within 4 weeks, representing more than 40% of all clade A newly sequenced isolates. The largest cluster at GC2 included 8 isolates, and was found in both NSW and QLD within 4 weeks in 2018, suggesting a likely outbreak. Therefore, clade A has the potential to cause small-scale outbreaks. This was supported by a previous local outbreak in QLD caused by phage type 26 [22], which belongs to clade A. Clade A isolates were historically found in samples related to chickens, eggs, dogs and barramundi in QLD [15]. Therefore, the environmental reservoir of clade A is likely to be diverse. Investigations of the food/environmental sources of such small clusters in the future would help confirm such potential small-scale outbreaks as well as prevent large-scale clade A outbreaks. By contrast, no GC2 clusters were observed for clade C, with only one cluster of two isolates found at GC5 in QLD. Thus, clade C mainly causes sporadic infections in QLD in Australia.

### Most Australian clade B isolates were likely to have been imported

Clade B is a global epidemic clade, which includes at least 10 lineages as defined in our previous study [17]. Seven lineages were observed in Australia, with MGT4-CC1 as the dominant lineage (>80%), followed by MGT4-CC30. MGT4-CC1 is the global epidemic lineage, which corresponds to the global epidemic clade/lineage in Feasey *et al*. [2] and Li *et al*. studies [44], and the HC100_87 cluster by Achtman *et al*. [16]. The global epidemic lineage (MGT4-CC1) included phage type 4 (PT4) strains [44] and is prevalent in all continents except North America [17]. Lineage MGT4-CC30 was mainly observed in Europe [17]. Because of the large number of isolates from the NSW outbreak cluster, MGT4-CC30 was ranked as the second-largest lineage of Australian *S*. Enteritidis. Excluding the outbreak isolates, the European lineage MGT4-CC30 is rare in Australia with only six isolates. Interestingly, three isolates obtained from extra-intestinal sources (blood, wound and ankle fluid) belonged to the two African MDR and invasive lineages (represented by MGT3-CC10 and MGT3-CC15) [2].

Australian clade B isolates from NSW and QLD were found to be frequently linked to isolates from other countries using higher resolution typing levels (MTG4 or above and GCs at MGT9 level). In the MGTdb, although there were a limited number of genomes from Asian countries, three out of five top MGT5-STs (all belonging to MGT4-CC1 lineage) in Australia were also top STs in Asia, suggesting that the Australian *S*. Enteritidis may be epidemiologically linked with Asian countries (**Table 2**). By GCs, 58.1% and 82.2% of Australian clade B isolates belonged to international clusters at GC5 and GC10, respectively (**Figure S2**). These findings were supported by data from the annual reports of OzFoodNet, 75% to 95% of *S*. Enteritidis infections were travel related from 2005 to 2011 and more than 80% of travel related *S*. Enteritidis infections were acquired from South-East Asia (mainly Bali, Thailand, and Malaysia) [18-20, 43], which are popular Australian tourist destinations. However, clade B isolates were also likely to be imported from other continents. For example, the Europe and North America prevalent lineage (MGT4-CC13) was also observed in Australia although less frequently. Overall, by comparison with international isolates using MGT, the majority of Australian clade B genomes were mainly linked to isolates from Southeast Asia followed by Europe.

Apart from international travel related infections, there were also potential small outbreaks or transmissions within Australia. At MGT9-ST, GC1, and GC2 levels, 16.5%, 27.5%, 34.7% of Australian clade B isolates from NSW and QLD were grouped together in MGT9 ST or GC clusters and were isolated within a 4-week window. These small clusters may be caused by a common infection source overseas; local secondary transmissions or consumption of the same imported contaminated food. The transmission from local farms to humans is less likely considering *S*. Enteritidis remains uncommon in Australian poultry industry [45, 46]. These findings highlight the need for further investigations as travel history or food exposure metadata which was not available in this dataset could help to better understand the transmission pathways in small closely related clusters and to monitor possible local sources of *S*. Enteritidis causing outbreaks.

### Imported Clade B can establish itself locally and cause outbreaks in Australia

*S*. Enteritidis was not usually considered to be a major cause of local outbreaks in Australia [21]. The egg-associated NSW outbreak caused by *S*. Enteritidis in 2018, is the first large scale local *S*. Enteritidis outbreak recorded in Australia [23, 24]. This outbreak caused more than 200 cases and spread to other Australian states (e.g. QLD and Victoria) and New Zealand [23, 24]. This study analysed 68 isolates of the outbreak using the MGTdb. We found three isolates from UK and Ireland were closely related (belonging to the same GC5 type) but were unlikely to be directly linked to the Australian outbreak because the three European isolates were collected from 2012 to 2015. By Bayesian analysis, the MRCA of the outbreak isolates was around 2015 (95% CI 2013-2017). The Australian isolates and international isolates (from the UK) had an MRCA around 2011 (95% CI 2008-2012) suggesting that this outbreak strain may have been imported into Australia around or after 2011, established locally and then caused a large outbreak.

These findings highlight the potential for travel-related or imported strains to colonise local farms and cause large outbreaks, despite stringent biosecurity protection against *Salmonella* in Australia [47]. Therefore, despite the low prevalence of *S*. Enteritidis in Australian farms [21], controlling the spread of imported strains and routine surveillance of clade B *S*. Enteritidis in the food production chain is critical. The establishment of an imported *S*. Enteritidis strain in Australian farms suggests that the absence of *S*. Enteritidis in local farms previously was not due to the farm environment in Australia being unsuitable for *S*. Enteritidis to survive and grow.

Importantly, this outbreak belonged to a recently defined lineage MGT4-CC30 [17], which was predominantly observed in Europe and the Australian outbreak is the first large-scale outbreak recorded for this lineage globally. More attention should be paid to this lineage as it can potentially cause large outbreaks, especially in Europe.

### AMR *S*. Enteritidis in Australia

MDR *S*. Enteritidis has been highlighted as an emerging threat to human health [2, 48]. Clade A and C isolates had no AMR genes identified. This was similar to the earlier study by Graham *et al*. [15]. The rarity of AMR genes in the Australian clades A and C reflected the relatively strict strategies in agricultural usage of antibiotics in Australia [49, 50]. In contrast, 69.6% of Australian clade B isolates possessed at least one AMR gene, and 11.7% of clade B isolates were predicted to be MDR. Using STs at different MGT levels, 54.8% of the MDR isolates belonged to previously identified MDR associated MGT STs [17], with MGT4-ST54, MGT6-ST107 and MGT4-ST354 as the major MDR associated STs. Interestingly, all these STs belonged to MGT3-ST8, which was one of the dominant types in Asian countries such as China [51, 52], South Korea and Thailand. These findings further confirmed that clade B *S*. Enteritidis strains isolated in Australia were likely to have originated from other countries, likely from Asia, as part of the global spread of these MDR clones. Continuous surveillance of MDR *S*. Enteritidis is critical because MDR associated infections are an increasing threat [2, 53-55]. MGT STs are potential good candidates for the global unified tracking of MDR clones. With more genomes available from countries where there is more antibiotics overuse or misuse, more MDR associated MGT STs are expected to be identified and their origins identified.

Plasmids are the main mechanism for the acquisition of AMR genes [56]. IncX1 plasmids (IncX1_4 and IncX1_1) were the most common plasmid replicon types in Australian clade B genomes and more than 80% of isolates with these plasmid types harboured AMR genes. The predicted MDR rate in isolates with plasmid IncX1_4 was nearly 40%, similar to our previous findings that a significantly higher proportion of isolates with IncX1_4 carried MDR genes [17]. Graham *et al*. also found that IncX1 plasmid was common in clade B *S*. Enteritidis that carried AMR genes [15]. An IncX1 plasmid with 16 AMR genes had been reported in *Salmonella* Agona from Australia [57]. Therefore, more attention should be paid to isolates with plasmid replicon type IncX1 (especially IncX1_4), which may have a higher risk of MDR acquisition and transmission.

### The application of MGT to epidemiological surveillance of *S*. Enteritidis

Public health laboratories across Australia have initiated genome-based surveillance within the communicable disease genomics network (CDGN) and a national microbial genomics framework has been formulated [58]. The Australian national genomics framework has outlined the following key requirements for the integration of genomic surveillance in public health systems [59]: **1**). Establishment of standardised national microbial genomic surveillance approaches using unified nomenclatures. **2**). Development of standardised genomic analysis reporting outputs for public health surveillance and response. **3**). Rapid and sustainable data sharing between public health laboratories. **4)**. An open-source platform for data processing and data repository is required.

The global open MGTdb for *S*. Enteritidis is a promising tool that can cater for all of these requirements. **1**). MGT uses standardised and consolidated nomenclature (STs/CCs of MGT) to identify predefined clades/lineage of *S*. Enteritidis; depict the geographic and temporal epidemiology using MGT STs; identify important clones (e.g. the MDR associated MGT STs); investigate potential outbreak clusters using the scalable resolution GCs. Thus, MGT enables standardised national and international microbial genomic surveillance. **2**). MGT consists of multiple allele-by-allele comparison-based MLST schemes [25], which does not require the construction of a phylogenetic tree and is computationally rapid. By describing the relationships of isolates using the numeric STs and cluster types at different MGT levels, the data interpretation could be simple and fast. To further simplify the data interpretation, the online platform allows users to select and visualise the epidemiological distribution of STs/CCs at different MGT levels (https://mgtdb.unsw.edu.au/enteritidis/summaryReport) [60]. **3**). An open database for *S*. Enteritidis is available (https://mgtdb.unsw.edu.au/enteritidis/), which allows for unified typing among different laboratories and comparison with publicly available international genomes. In summary, the comparison of the genomic features between states and countries using MGT offered a more comprehensive picture to understand *S*. Enteritidis in Australia. MGT is a promising system for surveillance of S. Enteritidis in Australia, and is a good candidate to be integrated into the public health surveillance system.

### Challenges of using public genome data

There are two challenges related to the use of publicly available isolates that are relevant to this study. Firstly, genome sequence datasets from different continents can be biased with Asia, South America and Africa being underrepresented. This means that international clusters were only described if they happened to be found in Australia and a region with better coverage of WGS data (North America/Europe). Secondly, global genomic surveillance relies on the availability and accuracy of metadata. Exact collection dates could facilitate more accurate outbreak investigations and source of samples or types of infections (e.g. gastroenteritis or iNTS) would facilitate the risk assessment of imported *S*. Enteritidis.

## Conclusion

This analysis of a large representative set of *S*. Enteritidis genomes using the MGT scheme illustrated the scalable resolution of MGT typing and demonstrated that Australian *S*. Enteritidis belonged to clades A, B and C. Clade A and C were confirmed to be endemic in Australia and were not linked to international clusters. However, clade A may have caused small-scale (2 to 12 isolates) local outbreaks. Clade B isolates were dominated by the global epidemic lineage MGT4-CC1 and likely to be mostly travel-related. The locally acquired egg-associated clade B outbreak cluster was found belonging to a lineage (MGT4-CC30) prevalent in Europe. AMR genes were only observed in clade B genomes and were especially common in isolates with plasmid type IncX1. *S*. Enteritidis MGT provides a systematic and sharable nomenclature and platform that can be effectively applied to public health agencies for unified surveillance of *S*. Enteritidis globally. The application of *S*. Enteritidis MGT platform to the Australian *S*. Enteritidis population in two states of Australia showcased its value for high-resolution epidemiological surveillance in Australia and internationally.

## Abbreviation

MGT: multilevel genome typing
MGTdb: MGT database
iNTS: non-typhoidal invasive infections
QLD: Queensland
NSW: New South Wales
MDR: multidrug resistance
AMR: Antimicrobial resistance
WGS: whole genome sequencing
MLST: multilocus sequence typing
cgMLST: core genome multilocus sequence typing
ODCs: outbreak detection clusters
GCs: genomic cluster
STs: sequence types
CCs: clonal complexes
ESS: effective sample size
MRCA: most recent common ancestor
CI: confidential interval
MLVA: multilocus variable-number tandem repeat analysis
PT: phage type
CDGN: communicable disease genomics network.

## Authors’ contributions

L.L., M.P. and R.L. designed the study. Q.W., I.R, R.G, M.G, J.D and E.M. collected data. M.P., R.L., S.O., Q.W., M.T., A.J., V.S. and S.K. provided critical analysis and discussions, L.L. wrote the first draft and all authors contributed to the final manuscript.

## Competing interests

The authors declare that they have no competing interests.

## Acknowledgements

This work was funded by grants from the National Health and Medical Research Council of Australia (1129713 and 2011806). Lijuan Luo was supported by a UNSW scholarship (University International Postgraduate Award). The funders had no role in study design, data collection and interpretation, or the decision to submit the work for publication. The authors thank Duncan Smith and Robin Heron from UNSW Research Technology Services for technical assistance.

## Notes

https://mgtdb.unsw.edu.au/enteritidis/

### Competing Interest Statement

The authors have declared no competing interest.

### Summary of Updates

The abstract was revised. An Importance section was added. The Bioproject of genomes were added.

## References

1. Achtman M, Wain J, Weill FX, Nair S, Zhou Z et al. Multilocus sequence typing as a replacement for serotyping in Salmonella enterica. PLoS Pathog, 2012;8(6):e1002776.

2. Feasey NA, Hadfield J, Keddy KH, Dallman TJ, Jacobs J et al. Distinct Salmonella Enteritidis lineages associated with enterocolitis in high-income settings and invasive disease in low-income settings. Nat Genet, 2016;48(10):1211–1217.

3. Ferrari RG, Rosario DKA, Cunha-Neto A, Mano SB, Figueiredo EES et al. Worldwide Epidemiology of Salmonella Serovars in Animal-Based Foods: a Meta-analysis. Appl Environ Microbiol, 2019;85(14):e00591–00519.

4. Patrick ME, Adcock PM, Gomez TM, Altekruse SF, Holland BH et al. Salmonella enteritidis infections, United States, 1985-1999. Emerg Infect Dis, 2004;10(1):1–7.

5. Cogan TA, Humphrey TJ. The rise and fall of Salmonella Enteritidis in the UK. J Appl Microbiol, 2003;94 Suppl(s1):114S–119S.

6. Dewey-Mattia D, Manikonda K, Hall AJ, Wise ME, Crowe SJ. Surveillance for Foodborne Disease Outbreaks -United States, 2009-2015. MMWR Surveill Summ, 2018;67(10):1–11.

7. Mansour MN, Yaghi J, El Khoury A, Felten A, Mistou MY et al. Prediction of Salmonella serovars isolated from clinical and food matrices in Lebanon and genomic-based investigation focusing on Enteritidis serovar. Int J Food Microbiol, 2020;333:108831.

8. Betancor L, Yim L, Fookes M, Martinez A, Thomson NR et al. Genomic and phenotypic variation in epidemic-spanning Salmonella enterica serovar Enteritidis isolates. BMC Microbiol, 2009;9:237.

9. Pijnacker R, Dallman TJ, Tijsma ASL, Hawkins G, Larkin L et al. An international outbreak of Salmonella enterica serotype Enteritidis linked to eggs from Poland: a microbiological and epidemiological study. Lancet Infect Dis, 2019;19(7):778–786.

10. Parn T, Dahl V, Lienemann T, Perevoscikovs J, De Jong B. Multi-country outbreak of Salmonella enteritidis infection linked to the international ice hockey tournament. Epidemiol Infect, 2017;145(11):2221–2230.

11. Inns T, Ashton PM, Herrera-Leon S, Lighthill J, Foulkes S et al. Prospective use of whole genome sequencing (WGS) detected a multi-country outbreak of Salmonella Enteritidis. Epidemiol Infect, 2017;145(2):289–298.

12. Hormansdorfer S, Messelhausser U, Rampp A, Schonberger K, Dallman T et al. Re-evaluation of a 2014 multi-country European outbreak of Salmonella Enteritidis phage type 14b using recent epidemiological and molecular data. Euro Surveill, 2017;22(50):17–00196.

13. Authority EFS, Prevention ECfD, Control. Multi□country outbreak of Salmonella Enteritidis phage type 8, MLVA profile 2□9□7□3□2 and 2□9□6□3□2 infections. EFSA support publ, 2017;14(3):1188E.

14. Assessment JE-ERO. Multi□country outbreak of Salmonella Enteritidis infections linked to eggs, third update –6 February 2020. EFSA support publ, 2020;17(2):1799E.

15. Graham RMA, Hiley L, Rathnayake IU, Jennison AV. Comparative genomics identifies distinct lineages of S. Enteritidis from Queensland, Australia. PLoS One, 2018;13(1):e0191042.

16. Achtman M, Zhou Z, Alikhan NF, Tyne W, Parkhill J et al. Genomic diversity of Salmonella enterica -The UoWUCC 10K genomes project. Wellcome Open Res, 2020;5(223):223.

17. Luo L, Payne M, Kaur S, Hu D, Cheney L et al. Elucidation of global and national genomic epidemiology of Salmonella enterica serovar Enteritidis through multilevel genome typing. Microb Genom, 2021;7(7).

18. Group OW. Monitoring the incidence and causes of diseases potentially transmitted by food in Australia: Annual report of the OzFoodNet network, 2011. Commun Dis Intell Q Rep, 2015;39(2):E236–E264.

19. Group OW. Monitoring the incidence and causes of diseases potentially transmitted by food in Australia: annual report of the OzFoodNet network, 2010. Commun Dis Intell Q Rep, 2012;36(3):E213–E241.

20. Group OW. Monitoring the incidence and causes of diseases potentially transmitted by food in Australia: annual report of the OzFoodNet Network, 2008. Commun Dis Intell Q Rep, 2009;33(4):389–413.

21. Ford L, Moffatt CRM, Fearnley E, Miller M, Gregory J et al. The Epidemiology of Salmonella enterica Outbreaks in Australia, 2001–2016. Front Sustain Food Syst, 2018;2(86).

22. Moffatt CR, Musto J, Pingault N, Miller M, Stafford R et al. Salmonella Typhimurium and Outbreaks of Egg-Associated Disease in Australia, 2001 to 2011. Foodborne Pathog Dis, 2016;13(7):379–385.

23. FSANZ. 2019. Salmonella Enteritidis (SE) linked to eggs https://www.foodstandards.gov.au/consumer/safety/Pages/Salmonella-Enteritidis-linked-to-eggs.aspx.

24. Svahn AJ, Chang SL, Rockett RJ, Cliff OM, Wang Q et al. Genome-wide networks reveal emergence of epidemic strains of Salmonella Enteritidis. Int J Infect Dis, 2022;117:65–73.

25. Payne M, Kaur S, Wang Q, Hennessy D, Luo L et al. Multilevel genome typing: genomics-guided scalable resolution typing of microbial pathogens. Euro Surveill, 2020;25(20):1900519.

26. Alikhan NF, Zhou Z, Sergeant MJ, Achtman M. A genomic overview of the population structure of Salmonella. PLoS Genet, 2018;14(4):e1007261.

27. Wood DE, Salzberg SL. Kraken: ultrafast metagenomic sequence classification using exact alignments. Genome Biol, 2014;15(3):R46.

28. Gurevich A, Saveliev V, Vyahhi N, Tesler G. QUAST: quality assessment tool for genome assemblies. Bioinformatics, 2013;29(8):1072–1075.

29. Robertson J, Yoshida C, Kruczkiewicz P, Nadon C, Nichani A et al. Comprehensive assessment of the quality of Salmonella whole genome sequence data available in public sequence databases using the Salmonella in silico Typing Resource (SISTR). Microb Genom, 2018;4(2).

30. Souvorov A, Agarwala R, Lipman DJ. SKESA: strategic k-mer extension for scrupulous assemblies. Genome Biol, 2018;19(1):153.

31. Tableau (version. 9.1). J Med Libr Assoc, 2016;104(2):182–183.

32. Argimon S, Abudahab K, Goater RJE, Fedosejev A, Bhai J et al. Microreact: visualizing and sharing data for genomic epidemiology and phylogeography. Microb Genom, 2016;2(11):e000093.

33. Hu D, Liu B, Wang L, Reeves PR. Living Trees: High-Quality Reproducible and Reusable Construction of Bacterial Phylogenetic Trees. Molecular biology and evolution, 2020;37(2):563–575.

34. Suchard MA, Lemey P, Baele G, Ayres DL, Drummond AJ et al. Bayesian phylogenetic and phylodynamic data integration using BEAST 1.10. Virus Evol, 2018;4(1):vey016.

35. Rambaut A, Drummond AJ, Xie D, Baele G, Suchard MA. Posterior Summarization in Bayesian Phylogenetics Using Tracer 1.7. Syst Biol, 2018;67(5):901–904.

36. Kumar S, Stecher G, Li M, Knyaz C, Tamura K. MEGA X: Molecular Evolutionary Genetics Analysis across Computing Platforms. Molecular biology and evolution, 2018;35(6):1547–1549.

37. Rambaut A. FigTree v1. 4. 2012.

38. Feldgarden M, Brover V, Haft DH, Prasad AB, Slotta DJ et al. Validating the AMRFinder Tool and Resistance Gene Database by Using Antimicrobial Resistance Genotype-Phenotype Correlations in a Collection of Isolates. Antimicrob Agents Chemother, 2019;63(11):e00483–00419.

39. Feldgarden M, Brover V, Gonzalez-Escalona N, Frye JG, Haendiges J et al. AMRFinderPlus and the Reference Gene Catalog facilitate examination of the genomic links among antimicrobial resistance, stress response, and virulence. Sci Rep, 2021;11(1):12728.

40. Carattoli A, Zankari E, Garcia-Fernandez A, Voldby Larsen M, Lund O et al. In silico detection and typing of plasmids using PlasmidFinder and plasmid multilocus sequence typing. Antimicrob Agents Chemother, 2014;58(7):3895–3903.

41. Feil EJ, Li BC, Aanensen DM, Hanage WP, Spratt BG. eBURST: inferring patterns of evolutionary descent among clusters of related bacterial genotypes from multilocus sequence typing data. J Bacteriol, 2004;186(5):1518–1530.

42. Sintchenko V, Wang Q, Howard P, Ha CW, Kardamanidis K et al. Improving resolution of public health surveillance for human Salmonella enterica serovar Typhimurium infection: 3 years of prospective multiple-locus variable-number tandem-repeat analysis (MLVA). BMC Infect Dis, 2012;12(1):78.

43. Group OW. Burden and causes of foodborne disease in Australia: annual report of the OzFoodNet network, 2005. Commun Dis Intell Q Rep, 2006;30(3):278–300.

44. Li S, He Y, Mann DA, Deng X. Global spread of Salmonella Enteritidis via centralized sourcing and international trade of poultry breeding stocks. Nat Commun, 2021;12(1):5109.

45. Chousalkar KK, McWhorter A. Egg Production Systems and Salmonella in Australia. In: Ricke SC, Gast RK (editors). Producing Safe Eggs. San Diego: Academic Press; 2017. pp. 71–85.

46. Chousalkar K, Gast R, Martelli F, Pande V. Review of egg-related salmonellosis and reduction strategies in United States, Australia, United Kingdom and New Zealand. Crit Rev Microbiol, 2018;44(3):290–303.

47. Bull A, Crerar S, Beers MY. Australia’s Imported Food Program-a valuable source of information on micro-organisms in foods. Commun Dis Intell Q Rep, 2002;26(1):28–32.

48. Musicha P, Cornick JE, Bar-Zeev N, French N, Masesa C et al. Trends in antimicrobial resistance in bloodstream infection isolates at a large urban hospital in Malawi (1998–2016): a surveillance study. Lancet Infect Dis, 2017;17(10):1042–1052.

49. Cox JM, Brook MD, Woolcock JB. Sensitivity of Australian isolates of Salmonella enteritidis to nitrofurantoin and furazolidone. Vet Microbiol, 1996;49(3-4):305–308.

50. Abraham S, Groves MD, Trott DJ, Chapman TA, Turner B et al. Salmonella enterica isolated from infections in Australian livestock remain susceptible to critical antimicrobials. Int J Antimicrob Agents, 2014;43(2):126–130.

51. Jiang M, Zhu F, Yang C, Deng Y, Kwan PSL et al. Whole-Genome Analysis of Salmonella enterica Serovar Enteritidis Isolates in Outbreak Linked to Online Food Delivery, Shenzhen, China, 2018. Emerg Infect Dis, 2020;26(4):789–792.

52. Deng Y, Jiang M, Kwan PSL, Yang C, Chen Q et al. Integrated Whole-Genome Sequencing Infrastructure for Outbreak Detection and Source Tracing of Salmonella enterica Serotype Enteritidis. Foodborne Pathog Dis, 2021;18(8):582–589.

53. Aldrich C, Hartman H, Feasey N, Chattaway MA, Dekker D et al. Emergence of phylogenetically diverse and fluoroquinolone resistant Salmonella Enteritidis as a cause of invasive nontyphoidal Salmonella disease in Ghana. PLoS Negl Trop Dis, 2019;13(6):e0007485.

54. Rodriguez I, Rodicio MR, Guerra B, Hopkins KL. Potential international spread of multidrug-resistant invasive Salmonella enterica serovar enteritidis. Emerg Infect Dis, 2012;18(7):1173–1176.

55. Vidovic S, An R, Rendahl A. Molecular and Physiological Characterization of Fluoroquinolone-Highly Resistant Salmonella Enteritidis Strains. Front Microbiol, 2019;10:729.

56. Reygaert WC. An overview of the antimicrobial resistance mechanisms of bacteria. AIMS Microbiol, 2018;4(3):482–501.

57. Cummins ML, Sanderson-Smith M, Newton P, Carlile N, Phalen DN et al. Whole-Genome Sequence Analysis of an Extensively Drug-Resistant Salmonella enterica Serovar Agona Isolate from an Australian Silver Gull (Chroicocephalus novaehollandiae) Reveals the Acquisition of Multidrug Resistance Plasmids. mSphere, 2020;5(6):e00743–00720.

58. Ferdinand AS, Kelaher M, Lane CR, da Silva AG, Sherry NL et al. An implementation science approach to evaluating pathogen whole genome sequencing in public health. Genome Med, 2021;13(1):121.

59. Health AGDo. 2019. National Microbial Genomics Framework 2019 – 2022. https://www.health.gov.au/resources/publications/national-microbial-genomics-framework-2019-2022.

60. Kaur S, Payne M, Luo L, Octavia S, Tanaka MM et al. MGTdb: A web service and database for studying the global and local genomic epidemiology of bacterial pathogens. bioRxiv, 2022:2022.2006.2014.496187.

61. Zankari E, Hasman H, Cosentino S, Vestergaard M, Rasmussen S et al. Identification of acquired antimicrobial resistance genes. J Antimicrob Chemother, 2012;67(11):2640–2644.

