## Supplementary material for "Genomic epidemiology and multilevel genome typing of Australian *Salmonella enterica* serovar Enteritidis"

**Supplementary Materials:**

**Supplementary Text**

**Epidemiological characteristics of the publicly available *S.* Enteritidis in the MGT database**

The newly sequenced Australian *S.* Enteritidis genomes were compared to the updated *S.* Enteritidis MGT database with 40,390 publicly available genomes from 99 different countries. Note that publicly available genomes were predominantly from the UK (41.6%) and the USA (37.3%). There were 12 countries from six continents with more than 100 genomes each (**Figure S1a**). In the publicly available dataset, there were a total of 22,696 (56.2%) genomes confirmed to be collected from humans, 3.0% (683/22,696) of which were indicated to be collected from blood or cerebrospinal fluids, indicating invasive human infections (**Figure S1b**). The second most common collection source was avian and avian-related products like eggs, which represented 10.5% (4256/40,390) of publicly available genomes. There were 30,903 (76.5%) publicly available genomes with collection year information, which ranged from 1917 to 2021 ((**Figure S1b**). There were 20,680 genomes with both collection year and month (but not day) information and 1838 genomes with collection date information (**Table S1**).

**Using GCs to identify international clusters in different clades**

The newly sequenced genomes were compared with the global *S.* Enteritidis MGT database to identify international clusters. Singletons referred to clusters with only one isolate. The non-singleton clusters were further grouped into international (found in two or more countries) and national (or Australian) clusters. For each clade, clusters at MGT9-ST, GC1 (i.e. MGT9-CC), GC2, GC5 and GC10 were categorised into international, national and singleton groups, respectively.

With the increasing number of allele differences allowed in single linkage clustering, there was a decreasing number of singletons observed from MGT9-ST to GC10 (grey columns in **Figure S2**). For clade A, all clusters of more than two isolates each were national (only observed in Australia) from MGT9-ST to GC10, except for one cluster at GC10 being international (observed in both Australia and UK) (**Figure S2a**). By contrast, the clade B *S.* Enteritidis included 12, 33, 56, 93 and 94 international clusters at MGT9-ST, GC1, GC2, GC5 and GC10, respectively. International clusters accounted for 3.2%, 11.2%, 18.8%, 58.1% and 82.2% of the total 473 clade B newly sequenced genomes at MGT9-ST, GC1, GC2, GC5 and GC10, respectively (**Figure S2b**). No international clusters were observed for clade C.

**Using GCs to identify interstate clusters in different clades**

The newly sequenced Australian *S.* Enteritidis were classified according to their distributions between the states. The non-singleton clusters (>= 2 isolates each) that were found in both NSW and QLD were referred to as interstate clusters. The non-singleton clusters that were observed in only one state were referred to as within QLD or NSW clusters. For each clade, clusters at MGT9-ST, GC1 (or MGT9-CC), GC2, GC5 and GC10 were categorised into interstate, within QLD, within NSW and singleton groups, respectively (**Figure S3**).

With the increasing number of allele differences allowed for single-linkage clustering, there was a decreasing number of singletons observed from MGT9-ST to GC10 (grey columns in **Figure S3**). For clade A, there were 1, 2, 3, 2 and 2 interstate clusters at MGT9-ST, GC1, GC2, GC5 and GC10, respectively. Those interstate clusters accounted for 4.8%, 11.9%, 17.9%, 19.0% and 19.0% of the total 84 clade A genomes at MGT9-ST, GC1, GC2, GC5 and GC10, respectively (**Figure S3a**). The clusters that were only found in QLD (within QLD clusters) were more frequent than those only found in NSW (within NSW clusters). For clade B, there were 2, 11, 14, 24 and 35 interstate clusters at MGT9-ST, GC1, GC2, GC5 and GC10, respectively (**Figure S3b**). Those interstate clusters accounted for 1.1%, 7.4%, 10.4%, 26.4% and 52.9% of the total 473 newly sequenced clade B genomes at MGT9-ST, GC1, GC2, GC5 and GC10, respectively. There was one large within NSW cluster observed from MGT9-ST (29 isolates) to GC10 (68 isolates) which is further described below. For clade C, there was only one within QLD cluster observed at GC5 and GC10, while the remainder are singletons.

### Evolutionary expansion of the Australian outbreak causing lineage MGT4-CC30

To determine the evolutionary origins of the local *S.* Enteritidis lineage, we identified its ST/CC at lower resolution MGT levels. At MGT5, the outbreak belonged to MGT5-ST810 while at MGT4 the outbreak isolates were assigned to the European prevalent lineage MGT4-CC30. To further examine the evolutionary dynamics of the outbreak and related isolates, a set of 144 isolates were randomly sampled from MGT5-ST810 and MGT4-CC30 for BEAST analysis. Strict molecular clock and Bayesian skyline model were evaluated as the optimal model with a mean ESS greater than 200. The most recent common ancestor (MRCA) of lineage MGT4-CC30 was estimated around 1942 (95% CI 1936-1951) based on the sampled isolates (**Figure S4a**). Lineage MGT4-CC30 includes two major sub-lineages, the MRCAs of which were 1959 (95% CI 1951-1967) and 1958 (95% CI 1952-1963), respectively (**Figure S4a**). By Bayesian skyline population expansion estimation, there was one large population expansion among the sampled isolates between 1980 to 1990, and another smaller population expansion between 2000 to 2010 (**Figure S4b)**.

### Supplementary Tables

#### Table S1. The collection time information of the genomes.

| | Australian | | | |
|---|---|---|---|---|
| | Newly sequenced | Publicly available | Non-Australian publicly available | Total |
| Day-Month-Year | 568 | | 1838 | 2406 |
| Month-Year | | 1 | 20,680 | 20,681 |
| Year | | 327 | 8385 | 8712 |
| Unknown | | | 9159 | 9159 |
| Total | 568 | 328 | 40,062 | 40,958 |

**Table S2. Number of isolates carrying different plasmid replicon types in different clades of Australian S. Enteritidis.**

| Plasmid replicon type | Clade A | Clade B | Clade C |
|---|---|---|---|
| IncFII(S)_1 | | 684 | |
| IncFIB(S)_1 | 2 | 680 | |
| IncX1_4 | | 159 | |
| Col156_1 [a] | 5 | 79 | |
| IncX1_1 | | 87 | |
| Col440I_1 | 1 | 38 | 1 |
| IncI1_1_Alpha | 8 | 31 | |
| ColRNAI_1 | 4 | 23 | |
| ColpVC_1 | 4 | 18 | |
| pSL483_1 | 1 | 10 | |
| IncI2_1_Delta | 5 | 1 | 1 |
| IncI_Gamma_1 | 1 | 4 | 1 |
| pESA2_1 | | 5 | |
| IncFII(pCTU2)_1_pCTU2 | | 5 | |
| IncX4_2 | | 4 | |
| Col(BS512)_1 | | 4 | |
| IncFII_1 | | 4 | |
| IncQ1_1 | | 3 | |
| IncX4_1 | | 3 | |
| IncFII(pHN7A8)_1_pHN7A8 | | 3 | |
| Others [b] | 2 | 16 | 1 |
| No. of isolates | 16 | 700 | 4 |

[a] Col156_1 can be observed twice in one isolate

[b] replicons with <3 isolates each

### Supplementary Figures

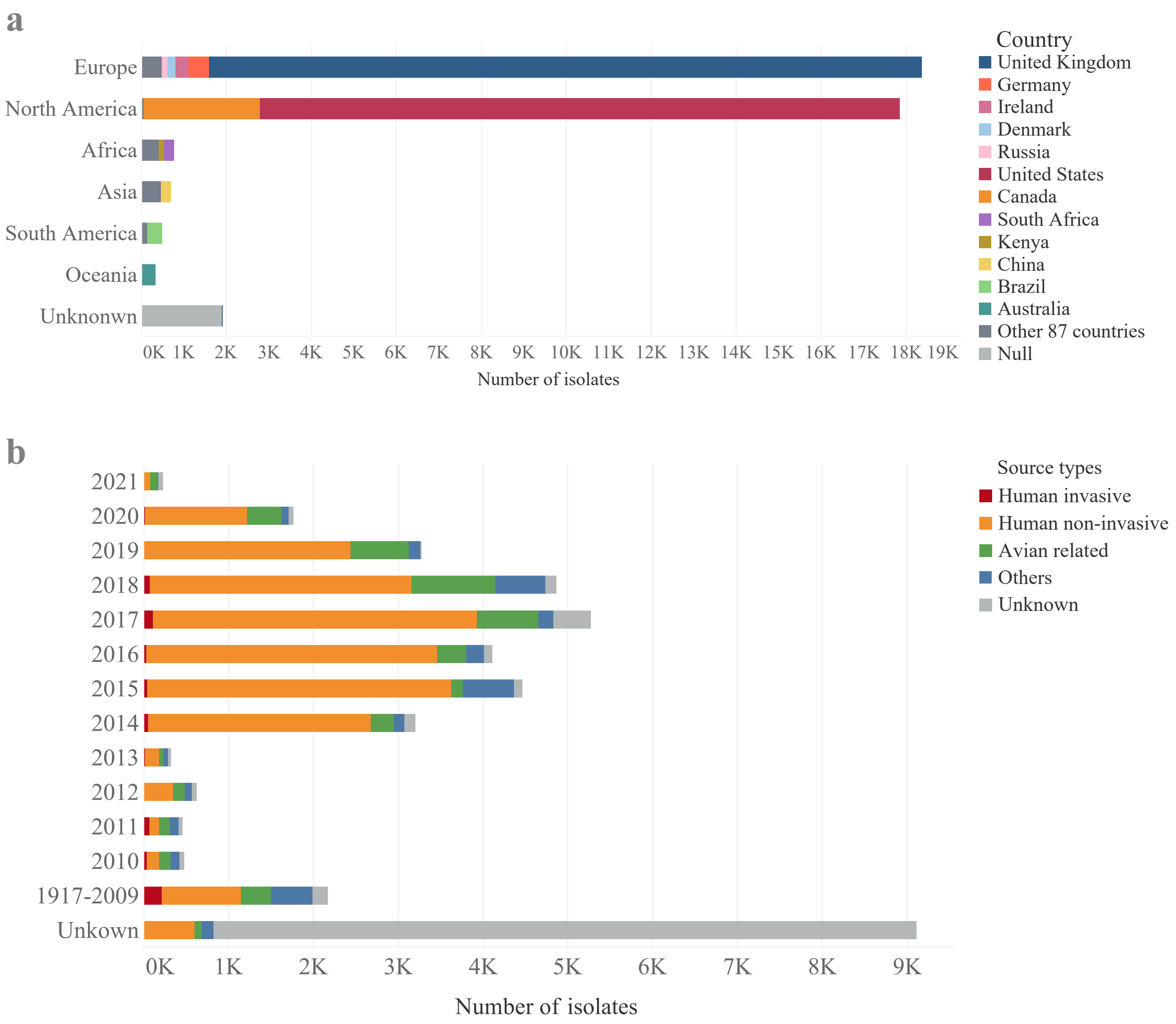


**Figure S1.** Epidemiological distribution of the publicly available genomes. **a**. Geographic distributions of the publicly available genomes. **b**. Collection years and source types of the publicly available genomes.

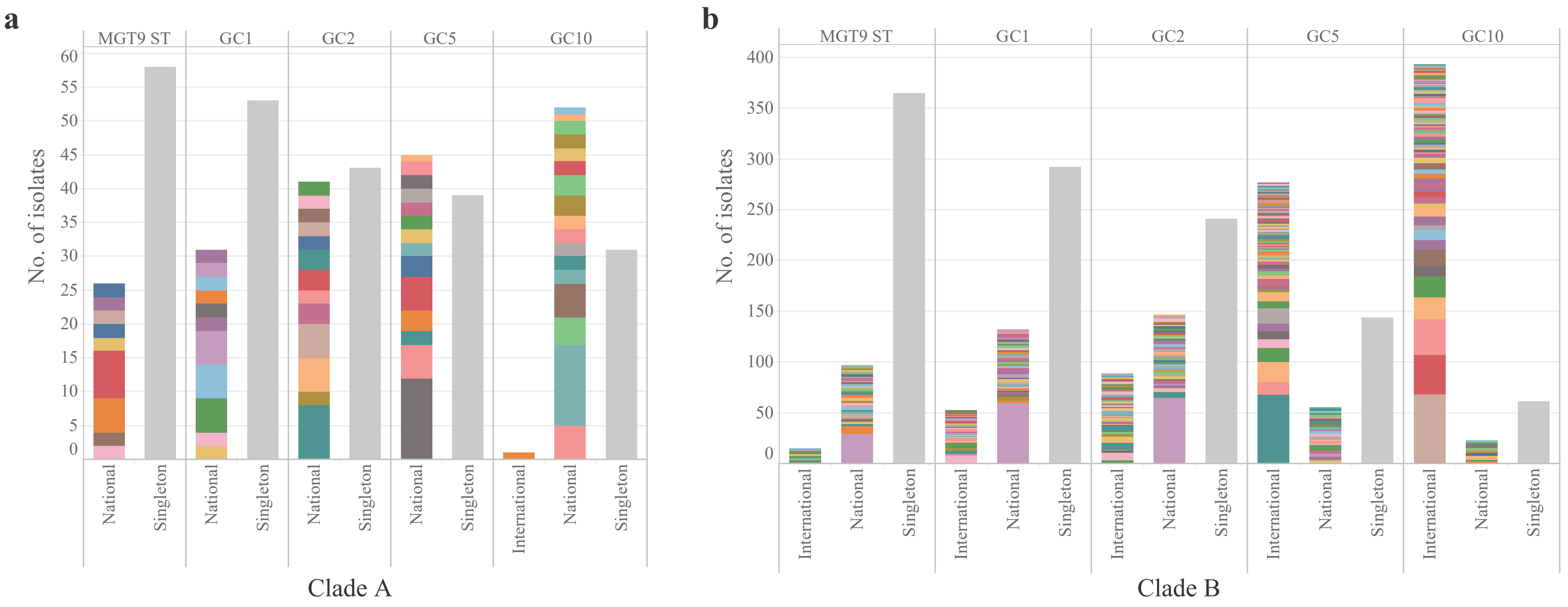


**Figure S2.** Geographical classification of GC clusters in clade A and B. Clusters at each GC level were categorised into three groups: Singletons with one isolate each (in grey colour); national clusters (>= 2 isolates each) that were only found in Australia; international clusters (>= 2 isolates each) that were also found in countries other than Australia. Note GC1 is the same as MGT9-CC. Different colours represented different single-linkage clusters.

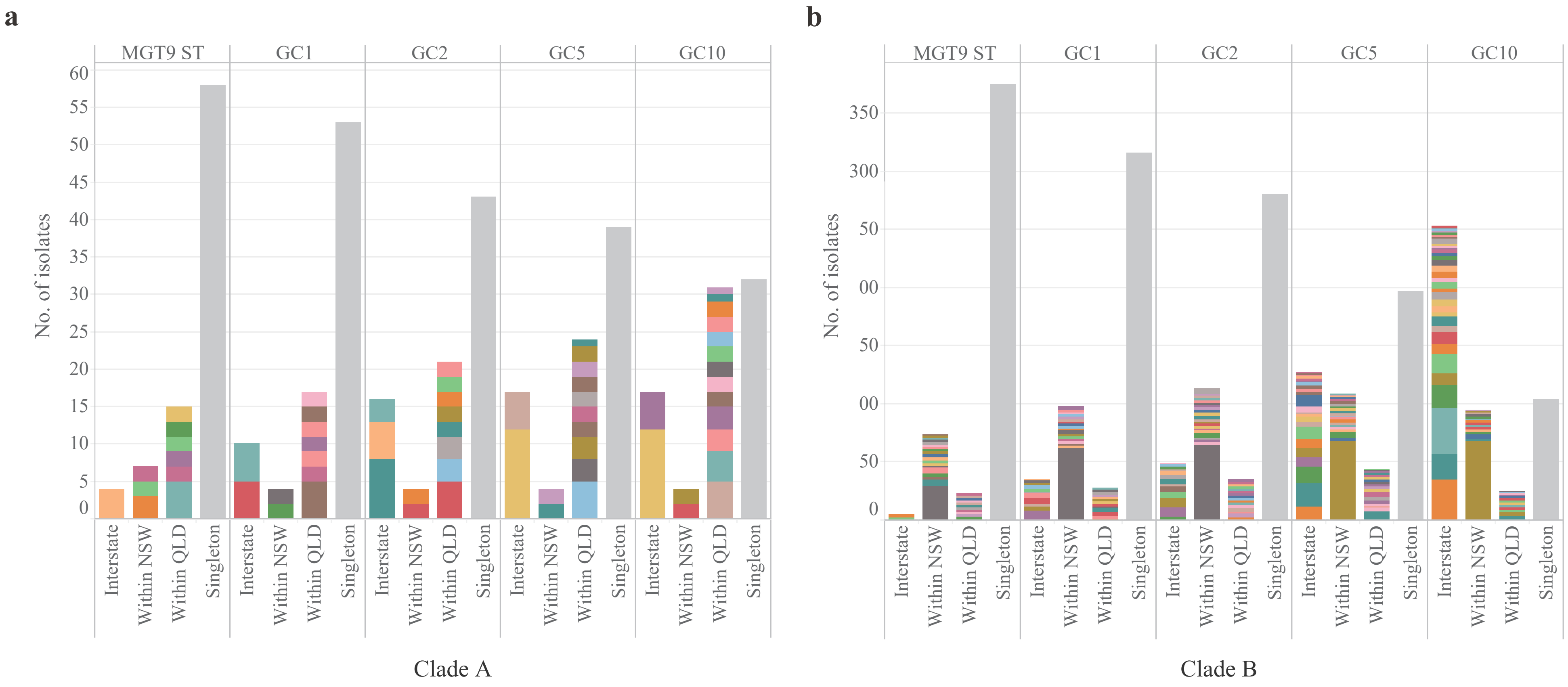


**Figure S3.** State distribution of GC clusters in clade A and B. Clusters at each GC level were categorised into four groups: singletons with only one isolate each in grey colour; interstate clusters (>= 2 isolates each) that were observed in both NSW and QLD; within NSW or QLD clusters (>= 2 isolates each) that were observed either in NSW or QLD. Note GC1 is the same as MGT9-CC. Different colours represented different single-linkage clusters.

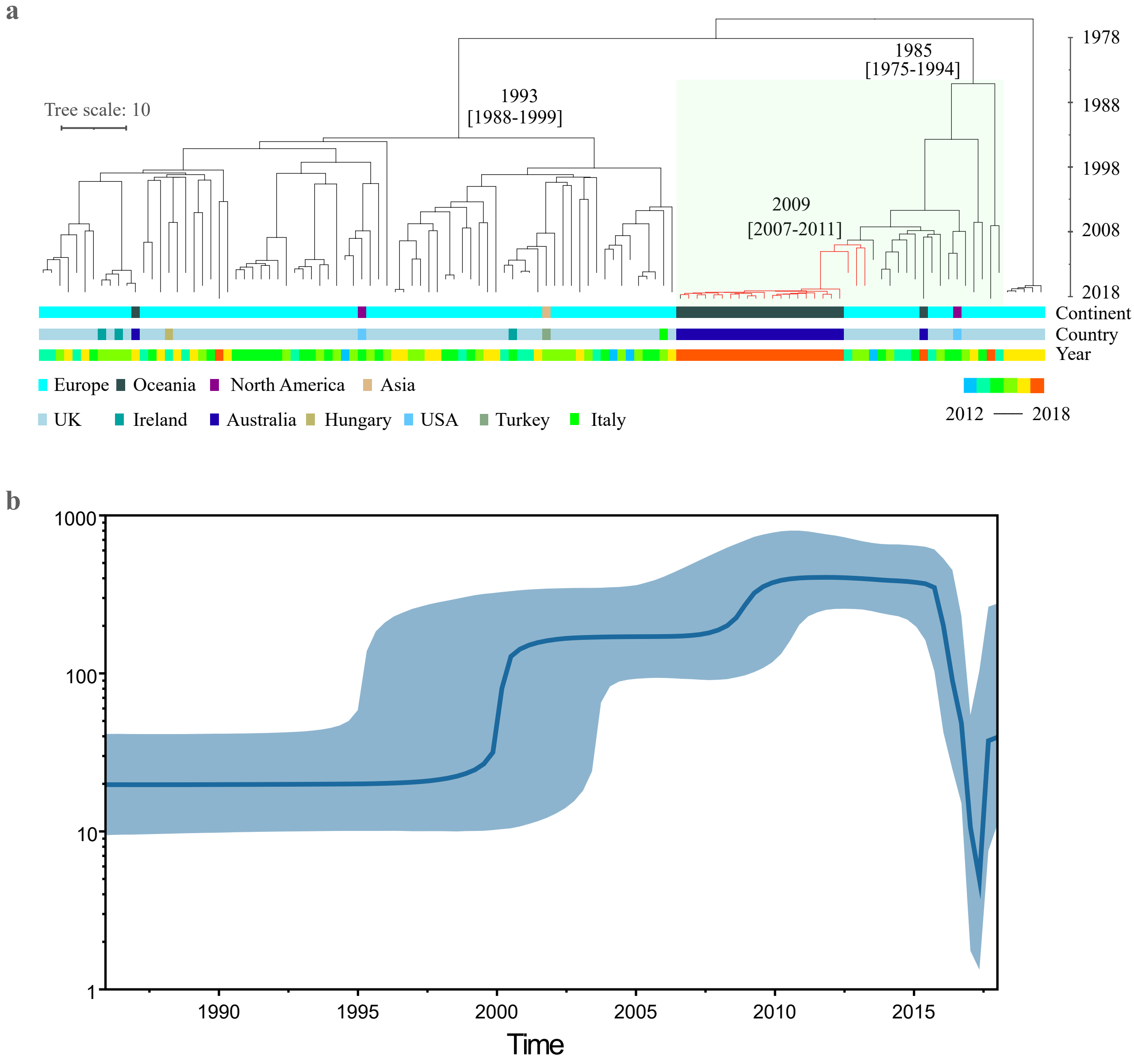


**Figure S4.** Phylodynamic analysis of the NSW outbreak-related isolates. The NSW outbreak belonged to the European epidemic lineage described by MGT4-CC30. A total of 144 isolates were randomly sampled for evolutionary analysis. **a**. Maximum likelihood tree of the outbreak-related isolates. Continent, country and collection year information of the sampled isolates were represented with different colours. This lineage includes two major sub-lineages, with one including the Australian outbreak isolates (with green background). The most recent common ancestor (MRCA) of this lineage and the two sub-lineages were indicated. **b**. The skyline population expansion of the lineage MGT4-CC30. There is one large population expansion between 1980 and 1990. The blue line represents the median posterior estimate of the effective population size. The blue area shows the upper and lower bounds of the 95% highest posterior density interval.

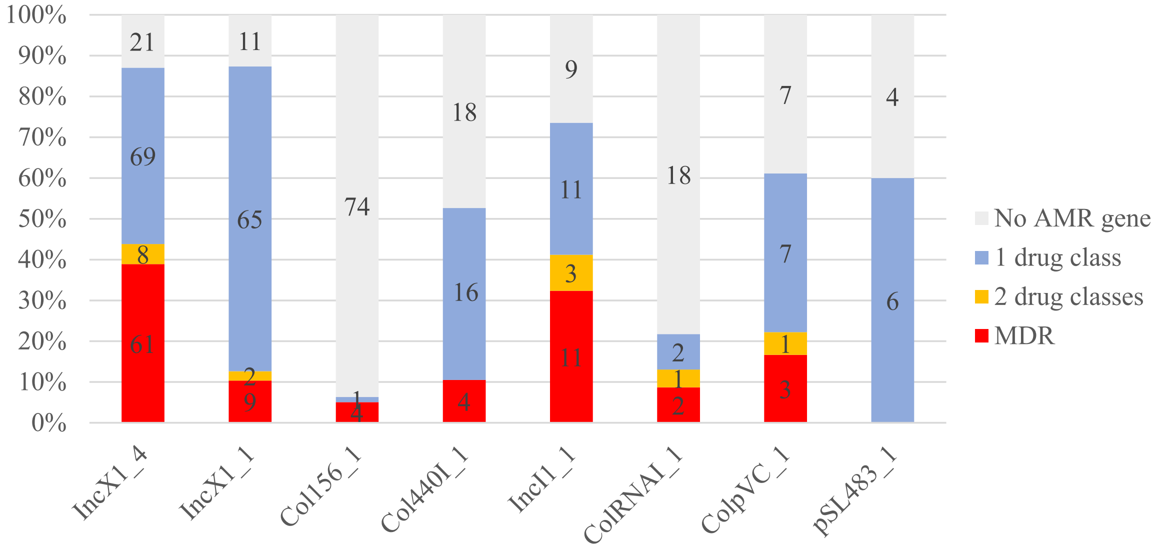


**Figure S5.** AMR gene distribution of isolates by plasmid replicon types. The numbers in the columns represented the number of isolates. Different colours represented the AMR gene distribution. MDR in red colour referred to isolates carrying AMR genes conferring resistance to three or more drug classes.
